# An investigation of PS-*b*-PEO polymersomes for the oral treatment and diagnosis of hyperammonemia

**DOI:** 10.1101/631630

**Authors:** Simon Matoori, Yinyin Bao, Aaron Schmidt, Eric J. Fischer, Rafael Ochoa-Sanchez, Mélanie Tremblay, Mariana Oliveira, Christopher F. Rose, Jean-Christophe Leroux

**Affiliations:** Institute of Pharmaceutical Sciences, Department of Chemistry and Applied Biosciences, ETH Zurich, Zurich, Switzerland; Institute for Chemical and Bioengineering, Department of Chemistry and Applied Biosciences, ETH Zurich, 8093, Zurich, Switzerland; Hepato-Neuro Laboratory, CRCHUM, Montréal, Québec, Canada

**Author notes:** John A. Paulson School of Engineering and Applied Sciences, Harvard University, Cambridge, MA 02138, USA. **Corresponding Author**: Prof. Dr. Jean-Christophe Leroux, ETH Zurich, HCI H 301, Vladimir-Prelog-Weg 3, 8093 Zurich, Switzerland.

**Keywords:** Polymersomes, poly(styrene)-*b*-poly(ethylene oxide), oral delivery, ammonia, hyperammonemia

## Abstract

Ammonia-scavenging transmembrane pH-gradient poly(styrene)-*b*-poly(ethylene oxide) polymersomes were investigated for the oral treatment and diagnosis of hyperammonemia, a condition associated with serious neurologic complications in patients with liver disease as well as in infants with urea cycle disorders. While these polymersomes were highly stable in simulated intestinal fluids at extreme bile salt and osmolality conditions, they unexpectedly did not reduce plasmatic ammonia levels in cirrhotic rats after oral dosing. Incubation in dietary fiber hydrogels mimicking the colonic environment suggested that the vesicles were probably destabilized during the dehydration of the intestinal chyme. Our findings question the relevance of commonly used simulated intestinal fluids for studying vesicular stability. With the encapsulation of a pH-sensitive dye in the polymersome core, the local pH increase upon ammonia influx could be exploited to assess the ammonia concentration in the plasma of healthy and cirrhotic rats as well as in other fluids. Due to its high sensitivity and selectivity, this novel polymersome-based assay could prove useful in the monitoring of hyperammonemic patients and in other applications such as drug screening tests.

**Graphical abstract:** 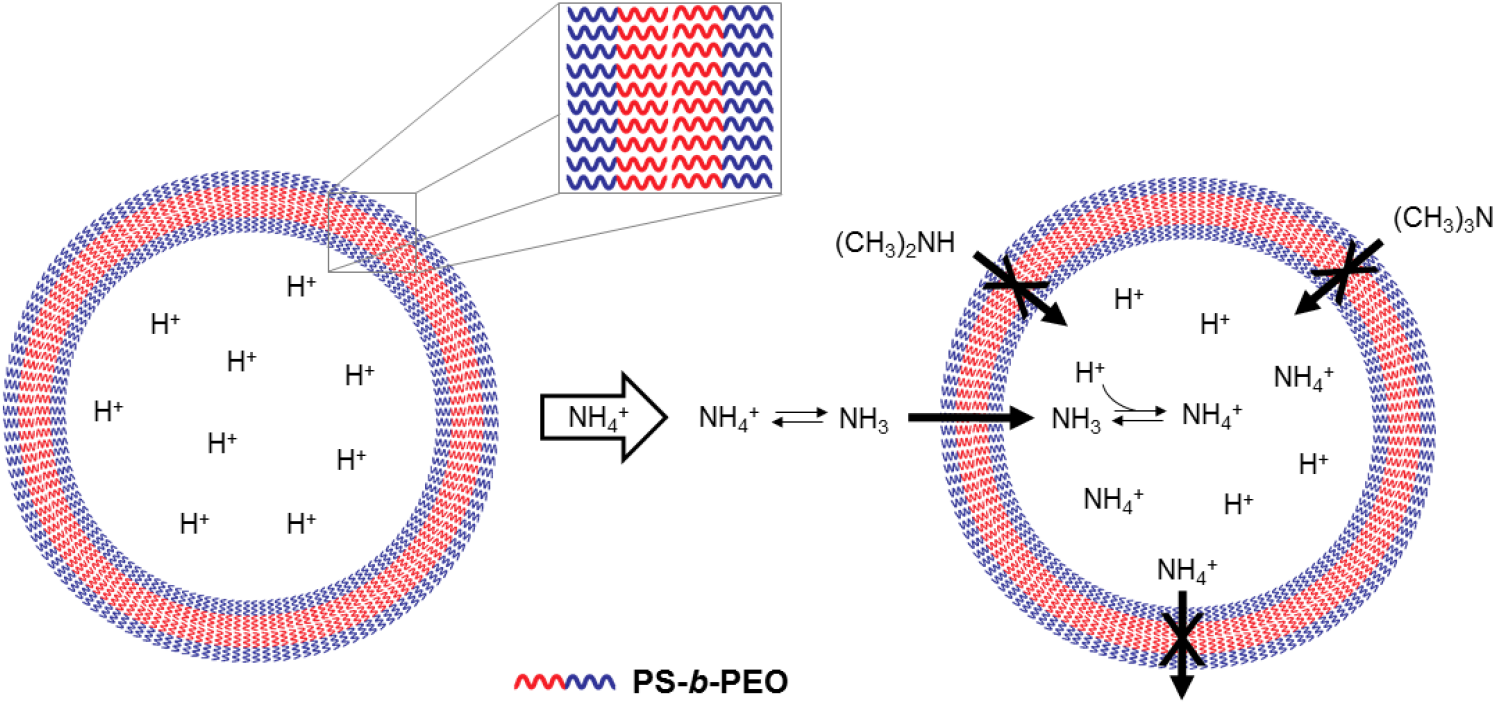

## 1. Introduction

Despite its important role in pH homeostasis and protein metabolism, the endogenous metabolite ammonia is associated with neurotoxic effects at pathologically elevated blood concentrations (hyperammonemia).^[1,2]^ As the liver is the main ammonia-removing organ, patients with inborn (*e.g.*, urea cycle disorders, UCD) or acquired liver disease (*e.g.*, liver cirrhosis, acute liver failure) often fail to efficiently clear ammonia.^[2]^ In consequence, ammonia accumulates in the systemic compartment, leading to neuropsychiatric symptoms in hyperammonemic patients (cognitive impairments, lethargy, hepatic coma) with a high risk of death.^[2]^ This syndrome termed hepatic encephalopathy (HE) affects up to 80% of patients suffering from liver cirrhosis.^[2]^ Current clinical practice guidelines for chronic HE patients primarily recommend treatments targeting the gut where urease-producing bacteria generate the main part of systemic ammonia.^[3]^ The first-line therapy lactulose is a laxative which leads to the expulsion of colonic ammonia (cathartic effect), and modulates the intestinal microbiome and pH.^[3]^ Despite its wide use in HE patients, the number of non-responding patients is high and its laxative properties negatively affect patient compliance.^[2]^ The second-line treatment rifaximin, a poorly absorbed antibiotic, inhibits the growth and metabolism of gut bacteria.^[3]^ However, the prolonged treatment with antibiotics bears the risk of inducing bacterial resistance.^[2]^

In both UCD and HE, elevated plasma ammonia concentrations cause cognitive deficits and impact on clinical outcome.^[4–8]^ Thus, ammonia levels are routinely measured for diagnosis, disease staging, prognosis, and monitoring during ammonia-lowering treatments.^[4–8]^ High accuracy and precision of quantification are essential in the context of hyperammonemia given the low plasma ammonia cut-off (50 µM in adults and 100 µM in newborns) and the correlation of peak ammonia levels (>200 µM in HE and UCD patients with acute hyperammonemia) with disease severity and clinical outcome.^[3–5,9,10]^ Yet, commonly used ammonia tests present numerous shortcomings. For instance, despite being the diagnostic gold standard, the L-glutamate dehydrogenase (GLDH)-based enzymatic assay has several interferences that are commonly found in liver disease patients (*e.g.*, hemolysis, icterus and hyperlipidemia).^[11]^ Moreover, the assay has limited throughput and is prone to timing errors as it relies on an enzymatic reaction. The alternative to the GLDH assay, the strip-based handheld device PocketChem BA,^[12]^ comparably underestimates ammonia levels and has similar drawbacks: low throughput (3 min for the measurement of one sample) and interference issues (*e.g.*, TMA and potentially other methylated ammonia analogs, Supplementary Figure 1).^[12]^

In former publications, we reported the use of transmembrane pH-gradient lipid vesicles (liposomes) to detoxify weakly basic drugs and endogenous metabolites such as ammonia.^[13–15]^ These substrates diffuse across the hydrophobic membrane in their deprotonated form and are trapped upon protonation in the acidic core of the vesicle. Applied as a peritoneal dialysis solution, such liposomes efficiently sequestered ammonia and decreased plasmatic ammonia levels in bile duct-ligated (BDL) rats, an established animal model of liver cirrhosis and HE.^[14]^ Although liposomal peritoneal dialysis is well adapted to the management of acute HA crises in a hospital setting, an oral treatment would be more suitable for the chronic treatment and prevention of HE. Therefore, this study aimed at investigating transmembrane pH-gradient vesicles for two other applications: the *in situ* detoxification of intestinal ammonia for the chronic treatment of HE and the quantification of ammonia in biological fluids for the diagnosis of hyperammonemia (Figures 1A & 1B).

**Figure 1.**
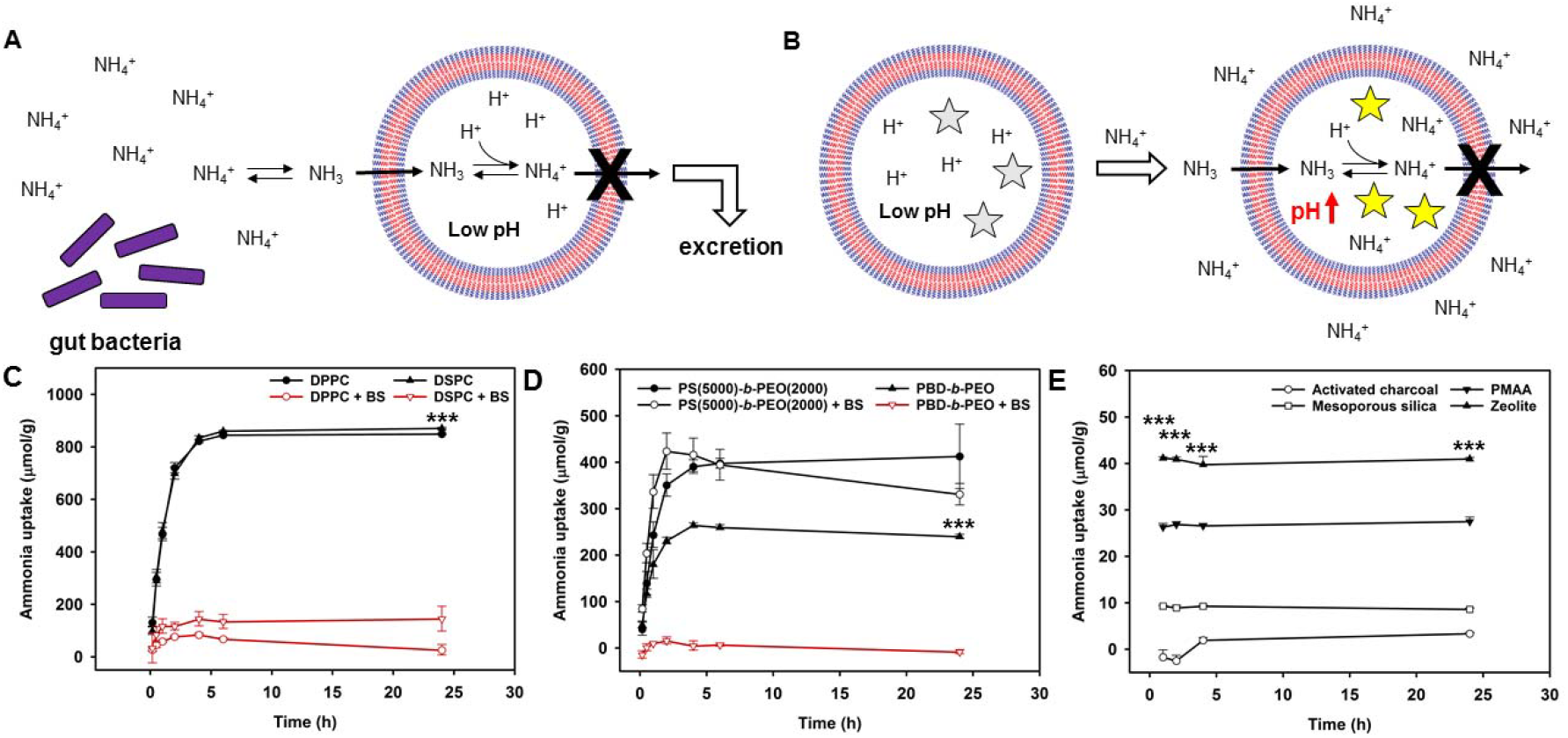
Ammonia sequestration by transmembrane pH-gradient lipo- and polymersomes at pH 6.8. Schematic depiction of intestinal ammonia sequestration (A) and ammonia sensing (B) using transmembrane pH-gradient polymersomes. Ammonia uptake of cholesterol-containing DPPC and DSPC liposomes at 1.75 mg/mL in isotonic buffer with or without bile salts (C). Ammonia uptake of polymersomes made of PBD(1500)-*b*-PEO(2000) or PS(5000)-*b*-PEO(2000) at 1.75 mg/mL in isotonic buffer with or without bile salts (D). Stars indicate significant difference in ammonia capture of vesicles in solutions with and without bile salts. Ammonia uptake of various microparticles (activated charcoal, PMAA, mesoporous silica, zeolite) at 30 mg/mL (E). Stars indicate significant difference of zeolite uptake from other microparticles. Buffer and bile salt composition: phosphate buffer 50 mM at pH 6.8; cholate, deoxycholate, taurocholate (25/25/0 mM for PBD(1500)-*b*-PEO(2000) polymersomes and liposomes; 30/30/30 mM for PS(5000)-*b*-PEO(2000) polymersomes); ammonia concentration 1.5 mM; temperature: 37°C. All results as means ± SD (n = 3). \*\*\**p* < 0.001.

## 2. Results and Discussion

### 2.1 Stability of liposomes and polymersomes in simulated GI fluids

The gastrointestinal (GI) tract is a harsh environment for vesicular structures due to its high bile salt concentrations, extreme osmolarity and pH values, and high enzymatic activity (*e.g.*, phospholipases),^[16]^ and may readily destabilize liposomes with phase transition temperatures (T_m_) below 37°C. Therefore, liposomes with high levels of cholesterol (45%) and relatively high T_m_ phospholipids (1,2-dipalmitoyl-sn-glycero-3-phosphocholine, *DPPC*, T_m_ = 41°C; 1,2-distearoyl-sn-glycero-3-phosphocholine, *DSPC*, T_m_ = 55°C) were first investigated. These liposomes had a mean size in the micrometer range (Supplementary Table S1) to minimize uptake by the GI mucosa. As shown in Figure 1C, the ammonia uptake reached up to 800 µmol NH_3_/g lipids in bile salt-free buffers but was completely abolished in the presence of physiologically^[17]^ relevant bile salt concentrations. More resistant, non-biodegradable micrometer-sized polymeric vesicles (Supplementary Table S1) were subsequently investigated to tackle this issue.^[18–21]^ They were prepared by emulsification from diblock copolymers of poly(ethylene oxide) (PEO) and poly(butadiene) (PBD) or poly(styrene) (PS) having glass phase transition temperatures (T_g_) of −92°C^[22]^ and 75°C,^[23]^ respectively. PBD-*b*-PEO polymersomes were readily destabilized by the bile salts (Figure 1D), and only the polymersomes made of the high T_g_ diblock copolymer PS-*b*-PEO preserved their ammonia capture capacity in simulated intestinal media (Figure 1D). The capture capacity of PS-*b*-PEO polymersomes was more than 15-fold higher than that of activated charcoal microparticles (AST-120, approx. 25 µmol NH_3_/g^[24]^), an ammonia-scavenging agent, which was reported to decrease plasmatic ammonia levels in BDL rats upon oral administration.^[24]^ The destabilization of liposomes and PBD-*b*-PEO polymersomes by surfactants was likely mediated by the insertion of the amphiphiles into the bilayer and the partitioning of the amphiphilic membrane constituents into the surfactant micelles.^[25–31]^ In contrast, the high glass transition temperature of PS yielded a much tougher membrane. Furthermore, different microparticles were also evaluated for ammonia uptake capacity *via* electrostatic binding or other non-covalent interactions (*e.g.*, aluminosilicate-based zeolites), but discarded due to their competition with physiologically relevant cations (Figure 1E, Supplementary Figure S2), and risk of inducing electrolyte imbalances *in vivo*.^[32–34]^

### 2.2 Optimization of the polymersome formulation

A library of bile salt-resistant PS-*b*-PEO(2000) polymersomes was subsequently prepared to select the optimal PS fragment length. The ammonia uptake was preserved in polymersomes with PS fragments between 2500 and 6000 over 24 h in a solution containing 90 mM bile salts (Figure 2A). This bile salt concentration was twice as high as extreme physiological ones,^[17]^ the PS-*b*-PEO polymersomes further maintained their ammonia capture capacity in bile salt-containing media with pronounced hypo- and hyperosmolalities (160 and 620 mOsmol/kg,^[35–37]^ Figure 2B). Moreover, the PS-*b*-PEO polymersomes captured ammonia in solutions containing bile salts and digestive enzymes (trypsin, chymotrypsin, lipase) or large excesses of cations (Supplementary Figures S3 & S4). To the best of our knowledge, this study is the first to provide evidence on the high stability of PS-*b*-PEO polymersomes with short PS fragments. One study reported that PS-*b*-PEO polymersomes composed of very long PS fragments (>25’000) were resistant to surfactants.^[38]^ As polymers with such large hydrophobic fragments are prone to aggregation, these polymersomes are associated with low yields in the vesicle preparation process (typically <2.5 mg/mL).^[19,38,39]^ In contrast, the emulsification-based preparation method used in this study in conjunction with relatively short PS fragments yielded highly concentrated PS-*b*-PEO polymersome dispersions with polymer concentrations exceeding 100 mg/mL. Furthermore, the investigation of different citric acid solutions as the inner phase pointed to an isotonic citric acid solution of 250 mM as the best compromise between ammonia capture capacity and citric acid exposure (Figure 2C). Finally, cryo-SEM analysis confirmed the vesicular morphology of the PS-*b*-PEO polymersomes (see arrows, Figure 2D).

**Figure 2.**
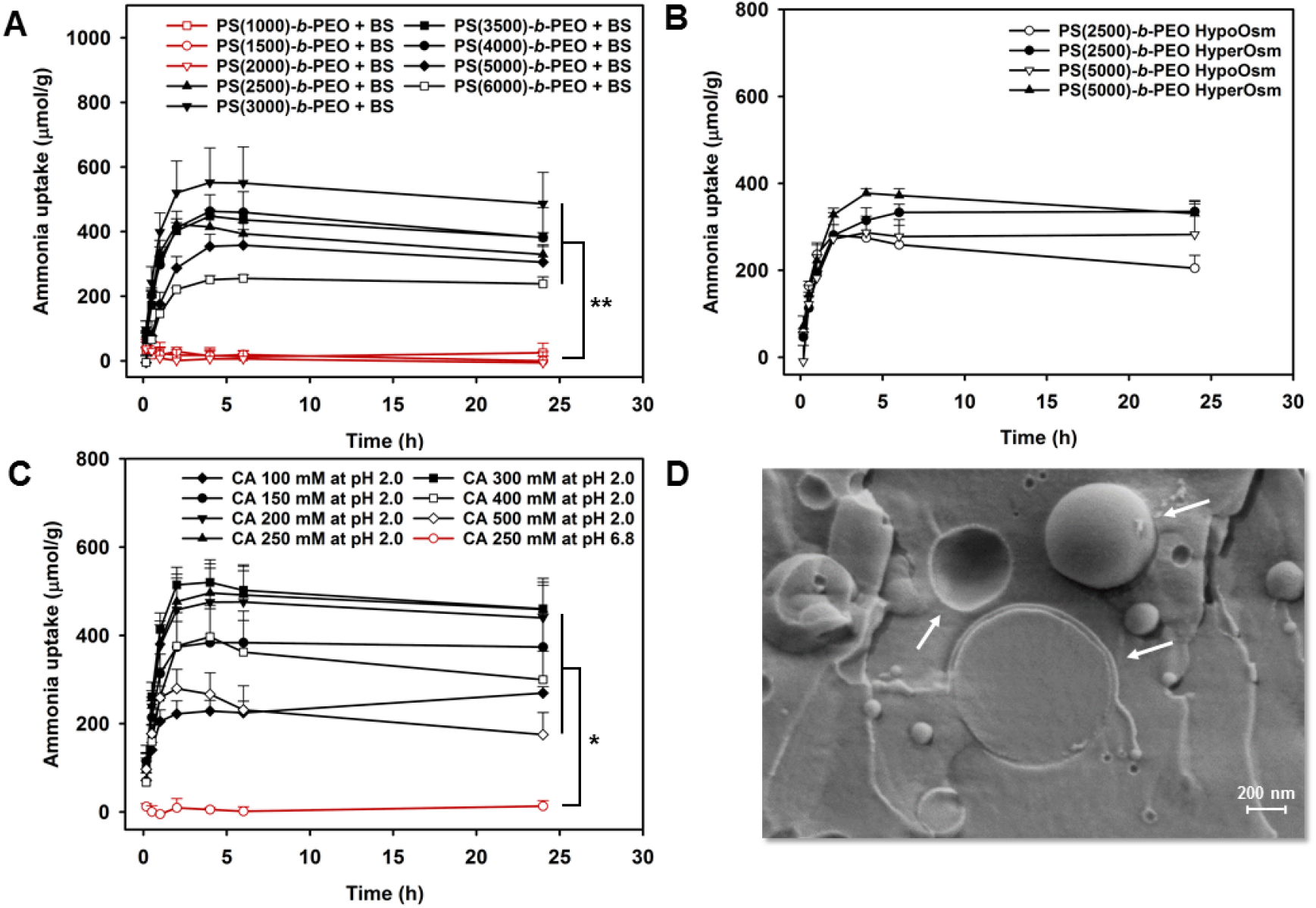
Ammonia uptake and characterization of PS-*b*-PEO polymersomes. Ammonia uptake of PS-*b*-PEO(2000) polymersomes with PS fragments between 1000 and 6000 in bile salt-containing buffer (A). Ammonia uptake of PS(2500)-*b*-PEO(2000) and PS(5000)-*b*-PEO(2000) polymersomes incubated in hypo- 160 mOsmol/kg) and hyperosmolal (620 mOsmol/kg) bile salt-containing buffer (B). Ammonia uptake of PS-*b*-PEO polymersomes with different citric acid concentrations in the core (100 to 500 mM) at pH 2.0 and control PS-*b*-PEO polymersomes without a transmembrane pH-gradient (core pH 6.8) in bile salt-containing buffer (C). Cryo-SEM image of PS(3300)-*b*-PEO(2000) polymersomes in isotonic citric acid solution 250 mM at pH 2.0 (D). Buffer and bile salt composition: phosphate buffer 50 mM at pH 6.8; cholate, deoxycholate, taurocholate (30/30/30 mM in all conditions except hypoosmolal conditions (25/25/0 mM)); ammonia concentration 1.5 mM; temperature: 37°C. All results as means ± SD (n = 3 (A, B), n = 3-8, C). \**p* < 0.05 and \*\**p* < 0.01.

### 2.3. Stability of polymersomes in dietary fiber hydrogels

To investigate the stability of the polymersomes under colon-mimicking conditions, a novel dietary fiber-based *in vitro* assay was established. PS-*b*-PEO(2000) polymersomes were incubated in dietary fiber (Metamucil^®^, psyllium husk) hydrogels at fiber concentrations reaching physiologically relevant stool water contents (approx. 50%, m/m), simulating the thickening of the feces in the colon due to water resorption. In other studies where liposomes were incorporated in hydrogels for drug delivery applications, the concentration of hydrogel-forming agents was typically much lower (approx. 2%, *m/m*) than the one used here (up to 50%, *m/m*).^[40–42]^ After 24 h of incubation, the polymersomes showed a fiber concentration-dependent decrease in ammonia capture capacity (Figure 3A). To investigate the destabilization mechanism in detail, the pH-dependent fluorescent dye 8-hydroxypyrene-1,3,6-trisulfonate (HPTS) was encapsulated in transmembrane pH-gradient polymersomes. With increasing fiber content, the fluorescence emission ratio (*i.e.*, emission at λ_em_ (510 nm) using pH-dependent λ_ex_ (413 nm) normalized to emission at the same λ_em_ using the isosbestic λ_ex_, 455 nm) of HPTS increased (Figure 3B), indicating a pH increase in the environment around the fluorescent dye (*i.e.*, loss of pH-gradient and/or efflux of dye).^[43]^ The disruption of the polymersome membrane was confirmed after encapsulating the dye/quencher pair HPTS/p-xylene-bis-pyridinium bromide (DPX) in the polymersome core, and observing an increase in HPTS fluorescence emission after hydrogel incubation (Figure 3C).^[44]^ The polysaccharide-based hydrogel-forming dietary fibers probably bound a large fraction of the free water molecules, resulting in a strong osmolality increase and a subsequent destabilization of the membrane. The comparably high water permeability of PS could allow an outflow of water in extremely hyperosmolal environments which may impair the polymersomes’ membrane integrity and result in the loss of the transmembrane pH-gradient.^[19,45]^ In addition, the ordered structure of the hydrogel might cause mechanical stress on the polymersome membrane, especially with regard to the large size of the polymersomes.^[46]^ A decrease in vesicle size might improve stability but bears the risk of increased uptake by M cells in the gut.^[47]^ While longer hydrophobic block lengths may make the polymersomes more resistant, their hydrophobicity could complicate the preparation of highly concentrated polymersome dispersions.^[19,38,39]^ Although PS-*b*-PEO polymersomes were stable in simulated intestinal fluids pushed to extreme bile salt and osmolality conditions, the instabilities detected in the dietary fiber assay cast doubt on the relevance of commonly used simulated intestinal fluids in the assessment of vesicular stability in the GI tract.^[30,48,49]^

**Figure 3.**
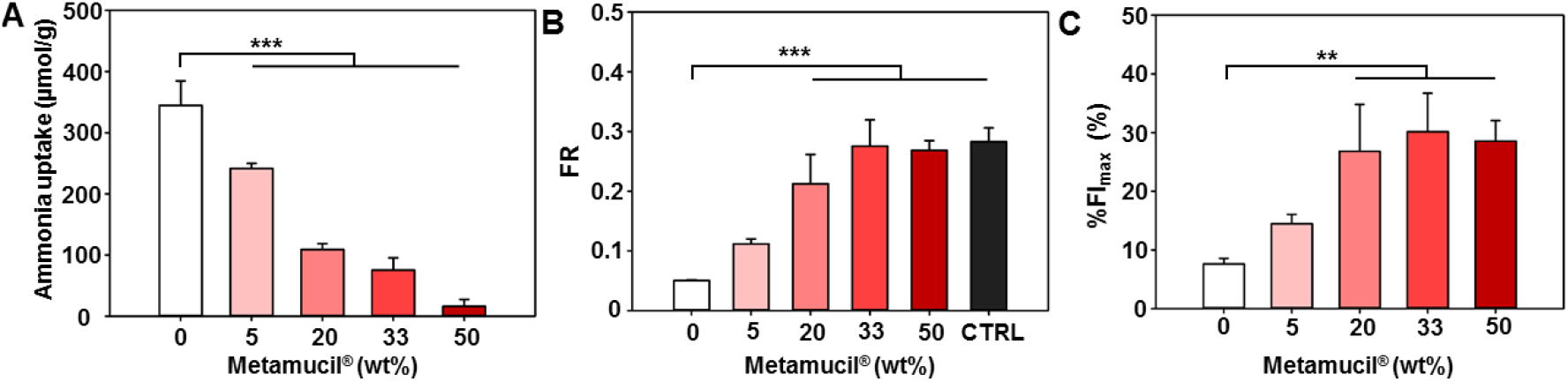
Stability of PS-*b*-PEO polymersomes in dietary fiber (Metamucil^®^)-based hydrogels. Ammonia uptake of transmembrane pH-gradient PS-*b*-PEO(2000) polymersomes after 24 h incubation in dietary fiber-based hydrogels at different fiber concentrations (A). Fluorescence emission ratio (FR, emission at λ_em_ (510 nm) using pH-dependent λ_ex_ (413 nm) normalized to emission at the same λ_em_ using the isosbestic λ_ex_, 455 nm) of HPTS-containing transmembrane pH-gradient PS-*b*-PEO polymersomes after 24 h incubation in dietary fiber-based hydrogels (B). Normalized fluorescence intensity (%FI_max_) of DPX/HPTS-containing polymersomes after 24 h incubation in dietary fiber-based hydrogels (C). PS fragment length: 3300-4500 g/mol; concentration of polymer (1 mg/mL, A) or dye (50 µM, B, 30 µM, C) in final dispersion; buffer composition: isotonic phosphate buffer at pH 6.8 (A-C); inner phase citric acid solution 250 mM at pH 2.0 (A, B, except for non-pH gradient control (pH 6.8) in B), isotonic phosphate buffer at pH 6.8 (C); temperature 37°C; n = 3. All results as means ± SD. \*\**p* < 0.01 and \*\*\**p* < 0.001.

### 2.4 Oral application of PS-*b*-PEO polymersomes

The PS-*b*-PEO polymersomes were orally administered to investigate their ammonia-scavenging capacity *in vivo*. In the light of the reduced stability of PS-*b*-PEO polymersomes in a simulated colon environment *in vitro*, laxative agents (PEG, sodium picosulfate) were orally applied to the BDL rats as an add-on to the polymersomes in order to hydrate the intestine. In contrast to PEG (1 g/kg), sodium picosulfate (25 mg/kg) significantly increased the stool water content compared with the water-only control over a period of 24 h (Figure 4A) in BDL rats. Hence, the efficacy of orally applied PS-*b*-PEO polymersomes was subsequently investigated in the presence of sodium picosulfate in BDL rats at 21 days after surgery with a seven-day twice-daily (1 g/kg per day) treatment (Figure 4B). The plasmatic ammonia levels were significantly increased in BDL rats at day 28 compared to non-BDL control rats. However, when comparing the two BDL groups, the plasma levels of the polymersome and the citric acid control group were not significantly different (Figure 4C). We hypothesize that the increase in stool hydration provided by the laxative was insufficient to prevent the destabilization of the polymersomes in the intestinal environment as they lose two thirds of their ammonia capture capacity in a dietary fiber hydrogel with a water content of 80% (m/m, *i.e.*, fiber content 20% (m/m), Figure 3A). The polymersome stability could potentially be improved by further increasing the laxative dose. However, excessive fluid and electrolyte losses could limit such a dose elevation. The present study shows that vesicles, which appear stable under highly challenging *in vitro* conditions, may still be destabilized in the GI tract, especially as the chyme becomes dehydrated. Therefore, more in-depth studies are warranted to investigate the stability of vesicular carriers in the GI tract, especially when they are used to deliver labile drugs such as insulin *via* the oral route.^[16]^

**Figure 4.**
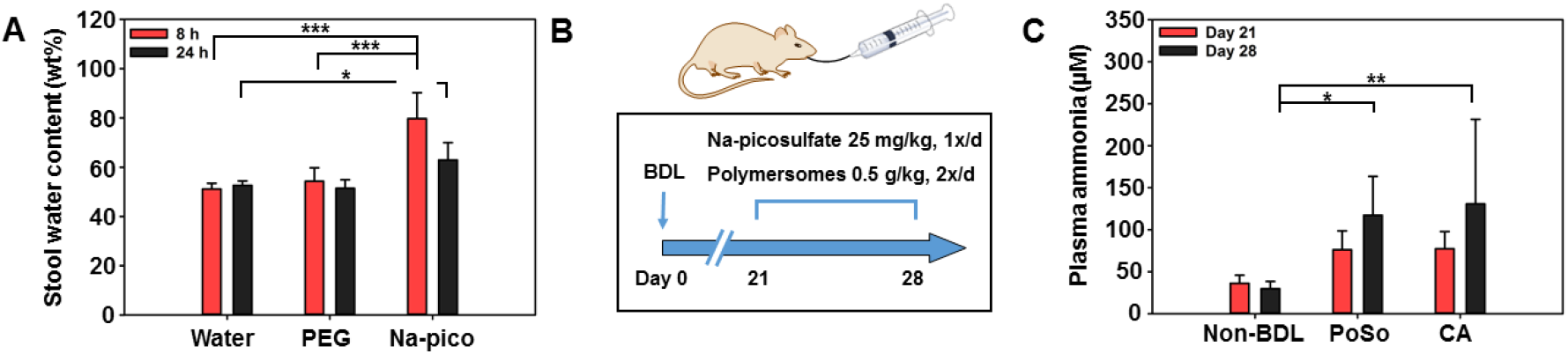
*In vivo* of PS-*b*-PEO polymersomes evaluation in BDL rats. Stool water content in male Sprague-Dawley rats 21 days after BDL surgery gavaged with water (5 mL/kg), PEG (3350) (PEG, 1 g/kg), or sodium picosulfate (25 mg/kg) for at least two days (n = 3-19 stool samples) (A). Study plan: three weeks after BDL surgery, two groups of cirrhotic rats (n = 16-17 per group) received sodium picosulfate (once daily, 25 mg/kg) and polymersome dispersion in citric acid solution (twice daily, 0.5 g PS-*b*-PEO(2000) polymer/kg; PS fragment length: 2800-5500 g/mol) or citric acid solution alone (twice daily 0.24 g citric acid/kg) by gavage for one week. A group of age-matched healthy control rats (n = 6) received citric acid solution (twice daily 0.24 g/kg) by gavage for one week (B). Plasma ammonia levels in BDL rats at 21 days (saphenous vein) and at sacrifice at 28 days (cardiac puncture) after BDL surgery (polymersome group n = 16, citric acid group n = 17) and in age-matched healthy controls (non-BDL control group n = 6, C). All results as means ± SD. \**p* < 0.05, \*\**p* < 0.01, and \*\*\**p* < 0.001.

### 2.5 Ammonia sensing with polymersomes

#### 2.5.1 Dye selection and characterization of polymersome-based ammonia assay

Notwithstanding, the transmembrane pH-gradient polymersomes may constitute a valuable diagnostic tool for HE and UCD, since adequate and affordable assays for the quantification of ammonia in biological solutions are still missing. Encapsulating the pH-sensitive dye HTPS in the polymersome core enabled the quantification of the increase in luminal pH upon ammonia influx and the determination of the extravesicular ammonia concentration (Figure 1B). As for the oral application, very high selectivity and stability of the vesicles are required for the diagnostic application. HPTS was selected due to its pH-responsive fluorescence excitation spectrum (Supplementary Figure S5) and its high hydrophilicity (three negative charges) which hinders its leakage. In addition, the isosbestic excitation wavelength allows for a correction of slight differences in dye concentration.

HPTS-containing PS-*b*-PEO polymersomes showed a linear ammonia sensing profile in the physiologically relevant range of 12.5 to 800 µM (Figure 5A). The concentrations of the calibration standards could be back-calculated with high accuracy and precision both below (non-hyperammonemic to moderately hyperammonemic region) and above 200 µM (highly hyperammonemic region) by splitting the linear regression curve or adjusting the volume fraction of the ammonia standard (Supplementary Figure S6, Supplementary Table S2). The linear range and coefficient of determination of the polymersome assay were comparable or possibly superior to the reported parameters for commercial or investigative ammonia tests such as the GLDH-based enzymatic assay and the PocketChem BA (Supplementary Figures S7 & S8, Supplementary Table S3). Under the investigated conditions, the lower limit of quantification was as low as 27 µM. Both a low and a high volume fraction of the ammonia standards showed a low time dependence of the fluorescence intensity ratio after 10 min at room temperature (Figure 5B) which points to the usefulness of the assay for high throughput screenings. Furthermore, PS-*b*-PEO polymersomes remained sensitive to ammonia after five months of storage in an isotonic phosphate buffer at pH 7.4 and 4°C (Supplementary Figure S9), highlighting the stability of the transmembrane pH gradient over time.

**Figure 5.**
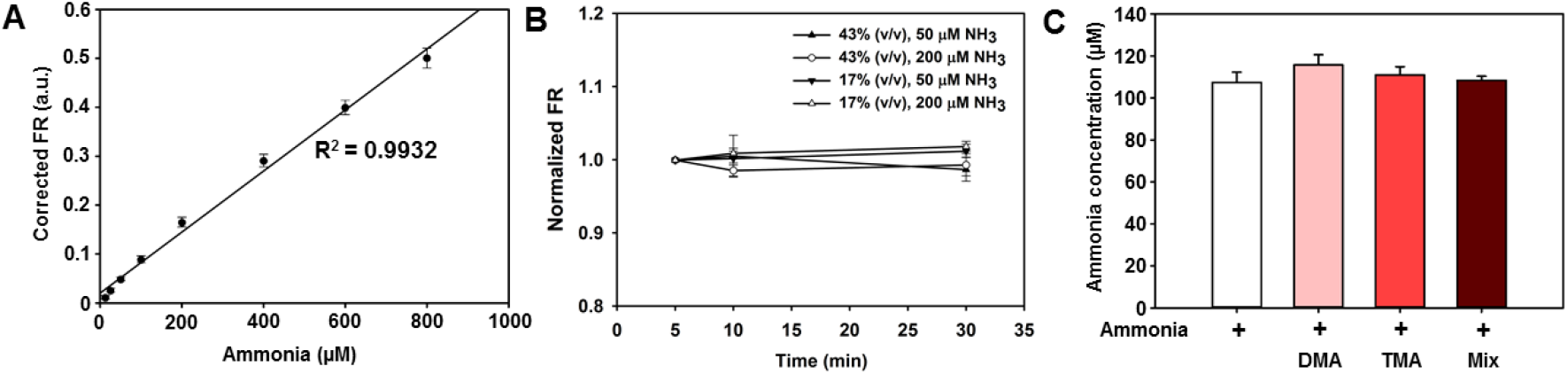
Ammonia sensing with HPTS-containing PS-*b*-PEO polymersomes. Blank-corrected fluorescence emission ratio (FR) of HPTS-loaded polymersomes at different ammonia concentrations (A). Normalized FR (FR at given time point normalized to FR at 5 min) of polymersomes exposed to 50 or 200 µM after 10 and 30 min incubation at room temperature. No statistically significant differences were observed between 10 min and 30 min for a given ammonia concentration and volume fraction of standard. Ammonia concentrations measured using the polymersome assay upon co-incubating ammonia (100 µM) with DMA (200 µM), TMA (100 µM), or a mixture (Mix) of drugs and endogenous compounds (levofloxacin, propranolol, glycine, alanine, dopamine, each at 500 µM) (C). No significant differences were observed between the ammonia concentrations determined in the absence and presence of the tested potentially interfering substances. Inner phase: isotonic citrate buffer 5 mM at pH 5.5; outer phase: isotonic phosphate buffer 50 mM at pH 7.4. HPTS concentration in final dispersion: 16 µM (A-C), 40 µM (B). Sample/standard volume fraction: 17% (A-C), 43% (B); PS fragment length: 2800-4500 g/mol; FR: emission at λ_em_ (510 nm) using pH-dependent λ_ex_ (413 nm) normalized to emission at the same λ_em_ using the isosbestic λ_ex_ (455 nm). All results as mean ± SD (n = 3).

#### 2.5.2 Selectivity of the polymersome assay

To assess the selectivity of the polymersome assay, PS-*b*-PEO polymersomes were co-incubated with amine-containing weakly basic substances (Supplementary Table S4). The di- and tri-methylated analogs of ammonia (dimethylamine, DMA, and trimethylamine, TMA, respectively) were chosen because of their structural similarity to ammonia and their physiological relevance.^[50,51]^ Even at concentrations exceeding physiological serum concentrations by more than 25-fold,^[50,51]^ DMA and TMA did not significantly influence the measured ammonia concentration (Figure 5C). Moreover, a mixture of drugs and metabolites (levofloxacin, propranolol, glycine, alanine, dopamine, each in five-fold molar excess over ammonia) did not interfere with the ammonia assay. PS-*b*-PEO polymersomes therefore selectively sequestered ammonia in the presence of weakly basic molecules that were shown to interfere with ammonia uptake in liposomes.^[15,52]^ The highly hydrophobic, rigid membrane of PS-*b*-PEO polymersomes probably hinders the diffusion of bulkier and more hydrophilic substrates than ammonia.

#### 2.5.3 Ammonia sensing in rat plasma

To evaluate the polymersome assay in plasma of healthy and BDL rats, the ammonia concentrations determined by the polymersome assay, the GLDH-based test, and the PocketChem BA were compared. In healthy rat plasma, the three tests performed similarly except for two hemolyzed samples in which the GLDH assay overestimated the ammonia levels compared with the other two tests (Figure 6A, Supplementary Figure S10). Hemolysis is a reported interference of the GLDH assay.^[11]^ In plasma of BDL rats, which exhibit typical pathological changes in biochemical laboratory parameters of liver cirrhosis (Supplementary Table S5),^[11,14]^ an interference in the polymersome assay was observed (Supplementary Figure S11), probably due to the high bilirubin levels. Indeed, bilirubin exhibits a broad fluorescence emission peak around 525 nm when excited at 455 nm.^[53]^ Lowering the sample volume fraction to 17% (*i.e.*, diluting the bilirubin) yielded similar results compared with the GLDH assay (Figure 6B). Alternatively, this interference might be circumvented by the replacement of HPTS with a pH-sensitive near-infrared dye which could further enable ammonia sensing in whole blood. In accordance with the literature, the PocketChem BA yielded significantly lower ammonia levels than the GLDH assay.^[12]^

**Figure 6.**
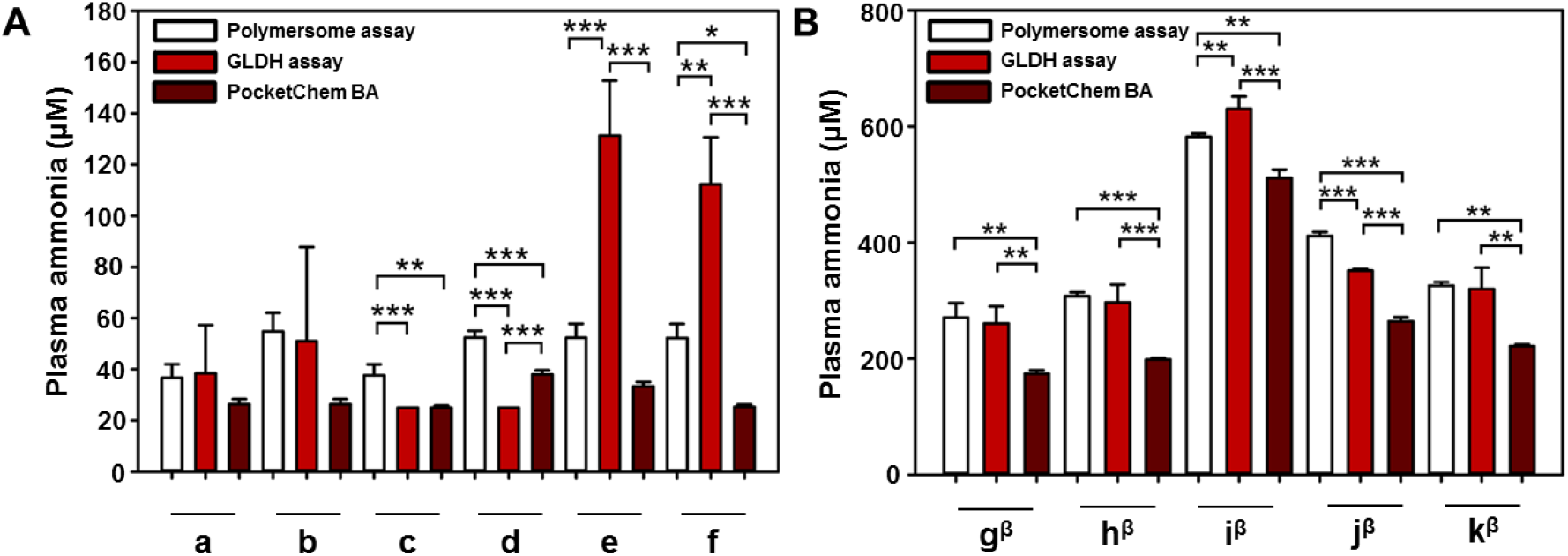
Ammonia sensing in rat plasma. Arterial plasma samples (cardiac puncture) from healthy (A) and BDL (β, four weeks after surgery, B) male Sprague-Dawley rats measured with the polymersome assay, GLDH-based ammonia assay, and PocketChem BA. Each letter (a-k) on the x-axis represents one animal. Inner phase: isotonic citrate buffer 5 mM at pH 5.5; outer phase: isotonic phosphate buffer 50 mM at pH 7.4. HPTS concentration in final dispersion: 40 µM (A), 16 µM (B). Sample/standard volume fraction: 43% (A), 17% (B); PS fragment length: 2800-4500 g/mol; FR: emission at λ_em_ (510 nm) using pH-dependent λ_ex_ (413 nm) normalized to emission at the same λ_em_ using the isosbestic λ_ex_ (455 nm). All results as mean ± SD (n = 3-10). \**p* < 0.05, \*\**p* < 0.01, and \*\*\**p* < 0.001.

#### 2.5.4 Ammonia sensing in other biological fluids

Interestingly, the polymersome-based ammonia assay was not limited to the determination of plasmatic ammonia levels in hyperammonemia. The polymersome assay was successfully used to quantify ammonia in various human body fluids such as saliva, urine, sweat, and semen upon lowering the sample volume fraction (Figure 7A). It was also capable of discriminating ammonia-spiked intestinal fluid samples (Figure 7B), which could be of interest in the establishment of a high-throughput screening for urease inhibitors. Furthermore, co-incubating the polymersomes with L-phenylalanine ammonia-lyase, which converts L-phenylalanine to ammonia and *trans*-cinnamic acid, enabled the quantification of phenylalanine (Supplementary Figure S12). The plasma levels of this amino acid are elevated in phenylketonuria, a hereditary disease caused by phenylalanine hydroxylase deficiency.^[54]^

**Figure 7.**
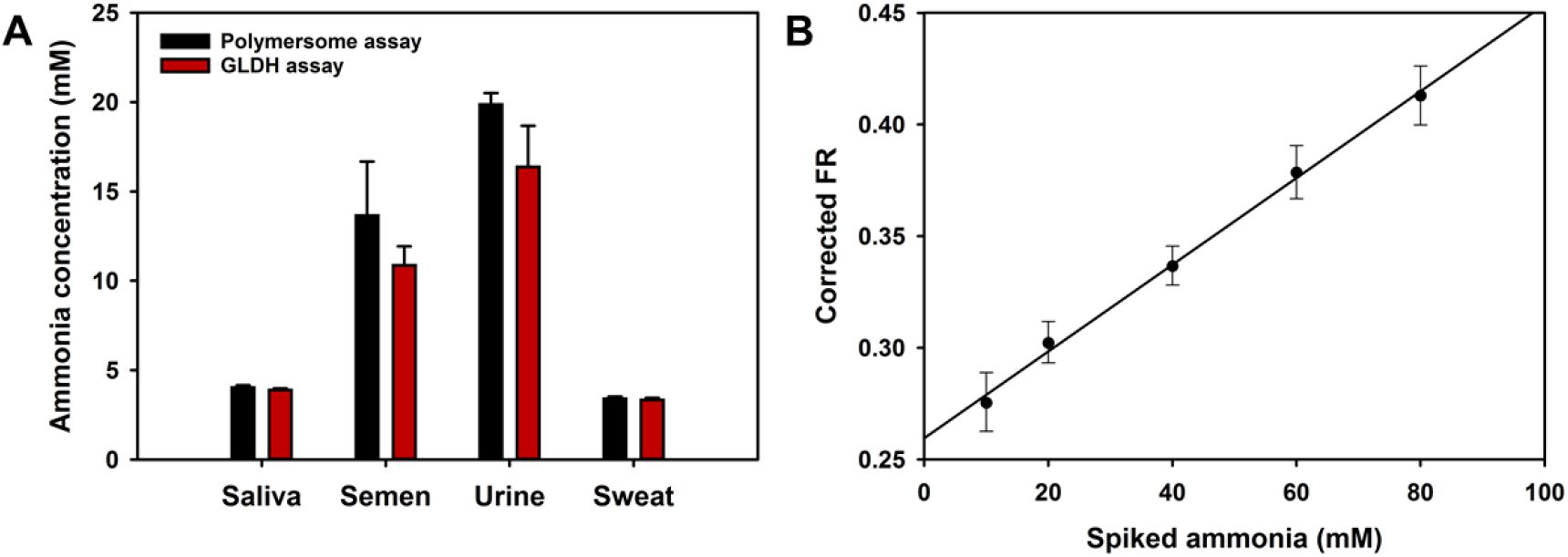
Ammonia sensing in various biological fluids. Ammonia concentration determined by the polymersome- and GLDH-based ammonia assays in human saliva, semen, urine, and sweat (A). No statistically significant differences were observed between the two ammonia assays. Baseline-corrected fluorescence emission ratio of HPTS-containing polymersomes in rat caecum fluid spiked with ammonium chloride (B). PS fragment length: 2800 (A), 4100 (B). Inner phase: isotonic citrate buffer 5 mM at pH 5.5, outer phase: isotonic phosphate buffer 50 mM at pH 7.4. HPTS concentration in final dispersion: 16 µM. Volume fraction of sample: 0.17% (semen, urine), 1.7% (saliva, sweat) (A), 0.5% (B). Corrected fluorescence emission ratio (FR): FR of blank subtracted from FR at given spiked ammonia concentration. All results as mean ± SD (n = 3).

## 3. Conclusion

In conclusion, PS-*b*-PEO polymersomes represent a promising sensitive and selective system for the monitoring of ammonia levels in plasma and various other body fluids, relevant for disease diagnosis and drug candidates screening. Despite being resistant to commonly used simulated intestinal fluids (bile salts, extreme osmolality conditions, digestive enzymes), these vesicles were destabilized under colon-mimicking conditions *in vitro* and in BDL rats *in vivo*. These results cast doubt on the relevance of such simulated intestinal fluids for studying the stability and predicting the release kinetics of vesicular delivery systems in the GI tract.

## 4. Experimental section

### 4.1 PS-*b*-PEO polymer synthesis and characterization

Poly(styrene)-*b*-poly(ethylene oxide) (PS-*b*-PEO(2000)) polymers were bought from Advanced Polymer Materials (Dorval, QC, Canada; Supplementary Table S6) or synthesized by atom transfer radical polymerization (ATRP, Supplementary Table S7).^[55]^ PEO(2000) monomethyl ether (Sigma-Aldrich Chemie, Buchs, Switzerland) was converted to an ATRP macroinitiator by reacting it with 2-bromopropionyl bromide (Sigma-Aldrich Chemie) in dry tetrahydrofuran (Acros Organics, New Jersey, NJ) and further used to polymerize styrene (Sigma-Aldrich Chemie) in bulk. Briefly, the ATRP macroinitiator (2.0 mmol) was loaded in a flame-dried Schlenk flask, along with copper bromide (CuBr, 3.0 mmol Alfa Aesar, Ward Hill, MA) and 4,4’-dinoyl-2,2’-dipyridyl (2.64 mmol, TCI, Tokyo, Japan) as the catalyst and ligand, respectively. The Schlenk flask was evacuated and refilled with argon for several cycles to remove oxygen. In a separate flask, styrene (100 mmol, 11.5 mL, Sigma-Aldrich Chemie) was deoxygenated by argon bubbling for 0.5 h, and then loaded in the Schlenk flask. The mixture was then heated at 115°C during 16 h and the solution containing the black product was dissolved in THF, filtered through a basic alumina column and precipitated twice in hexane. The precipitate was collected and dried under vacuum. The feeding molar ratio of [monomer]/[initiator] of 50 was used to achieve PS(4000)-*b*-PEO(2000). The PS/PEO composition was determined in deuterated acetone by ^1^H NMR (Bruker AV-400, Billerica, MA) at room temperature (Supplementary Figure S13).

For the molecular weight and dispersity (Đ) determination of the PS-*b*-PEO diblock co-polymers (Supplementary Table S7), the polymers were dissolved in THF at 2 mg/mL and characterized by gel-permeation chromatography with the same organic solvent (0.5 mL/min) as mobile phase equipped with two ViscoGEL columns (GMHHR-M, poly(styrene-*co*-divinylbenzene)) at 35°C and coupled to a refractive index detector (Viscotec GPCmax VE-2001 with Viscotek TDA 302 Triple Detector Array, Viscotek, Malvern Panalytical, Malvern, UK) using poly(methylmethacrylate) standards (2500 - 89300, PSS Polymer Standards Service, Mainz, Germany).

### 4.2 Polymersome preparation

PBD(2500)-*b*-PEO(1500) (number-averaged molecular weights, Đ 1.04, Polymer Source, Dorval, Canada) and PS-*b*-PEO polymersomes were produced using an oil-in-water (o/w) emulsification method. All polymers (60 mg) except PS(6000)-*b*-PEO(2000) were dissolved in 100 µL of dichloromethane (Sigma Aldrich Chemie). PS(6000)-*b*-PEO(2000) (20 mg) was dissolved in 100 µL of toluene (Sigma Aldrich Chemie). PS-*b*-PEO(2000) polymers of different PS fragment length were never pooled prior to polymersome preparation. The polymer solution was added dropwise to 1 mL citric acid (Sigma Aldrich Chemie) solution 250 mM at pH 2.0 at 300 mOsmol/kg (unless stated otherwise) under sonication in an ice bath using the following parameters: amplitude 70, cycle 0.75 (UP200H, 200W, 24 kHz equipped with sonotrode S1, Hielscher Ultrasonics, Teltow, Germany) for 3 min or amplitude 10 (3.1 mm sonotrode, Fisher Scientific Model 705 Sonic Dismembrator, 700W, 50/60 Hz, Fisher Scientific, Reinach, Switzerland) for 2 min. The organic solvent was evaporated with a rotary evaporator for at least 5 min at 40°C at 70 kPa.

In the scaled-up method, 1.84 g PS-*b*-PEO polymer (diblock copolymers from Supplementary Table S7; PS-*b*-PEO polymer batches with different PS fragment length were never pooled prior to polymersome preparation) were dissolved in 1.84 mL dichloromethane. The polymer solution was added dropwise to 16 mL citric acid solution 250 mM at pH 2.0 at 300 mOsmol/kg under sonication in an ice bath using the following parameters: initial amplitude 50 (6 mm sonotrode, Fisher Scientific Model 705 Sonic Dismembrator) during the addition of the oil phase (approx. 1 min) and subsequent sonication at amplitude 60 for another 3 min. The organic solvent was evaporated using a rotary evaporator for at least 10 min at 40 °C at 60 kPa. The polymersome batches were subsequently pooled. The concentration of dichloromethane in the pooled polymersome batches was further reduced by rotary evaporation at 40 °C and 22 kPa for 8 to 13 h. The residual dichloromethane content as determined by head-space gas chromatography (see section 4.3.2) ranged between 0.13 and 0.33 mg/mL.

The preparation of fluorescent PS-*b*-PEO polymersomes was conducted similarly to the low-volume protocol with a modified polymer amount (30 mg) and buffer composition. For the ammonia sensing experiments, an isotonic citric acid solution 5 mM at pH 5.5 containing 10 mM HPTS was chosen. For the dietary fiber hydrogel experiments (see section 4.6), an isotonic citric acid solution 250 mM at pH 2.0 containing HPTS 10 mM was used for the preparation of HTPS-containing transmembrane pH gradient polymersomes. As a control, non-pH gradient polymersomes containing an isotonic phosphate buffer 10 mM and HPTS 10 mM at pH 6.8 as the inner phase were prepared. For the preparation of HPTS/DPX-containing polymersomes, and an isotonic phosphate buffer 50 mM at pH 6.8 containing HPTS 10 mM and *p*-xylene-bis(*N*-pyridinium bromide) (DPX) 30 mM was selected.

The fluorescent polymersomes were purified using MidiTrap G-25 columns (GE Healthcare Europe, Freiburg i. Br., Germany) to remove the free dye and exchange the external phase with an isotonic sodium chloride-containing phosphate buffer (ammonia sensing: phosphate concentration 50 mM, pH 7.4; hydrogel experiments: phosphate concentration 10 mM, pH 6.8).^[44]^ In brief, after washing the column three times with the final buffer of choice, 200 µL of HPTS-containing polymersome dispersion and 800 µL of buffer solution were added to the column. The purified fluorescent polymersomes were eluted with 600 µL of buffer and stored at 4 °C protected from light.

### 4.3 Characterization of the polymersomes

#### 4.3.1 Concentration of polymer in the polymersome dispersions

The polymer concentration of PS-*b*-PEO was determined by spectrophotometry. The dispersion was diluted in *N,N*-dimethylformamide (final polymersome dispersion concentration 5% (v/v) and 2.5% (v/v) for 1 mL and for 16 mL set-up, respectively) by absorbance measurements at 271 nm (Tecan Infinite 200 Pro, Maennedorf, Switzerland). The polymer concentration was calculated using a PS-*b*-PEO standard curve in the same citric acid solution / *N,N*-dimethylformamide mixture.

The polymer concentration of PBD-*b*-PEO in the polymersome dispersion was determined by gel-permeation chromatography. After lyophilization, the polymersomes were dissolved in *N,N*-dimethylformamide and analyzed using gel-permeation chromatography with the same organic solvent (0.5 mL/min) as mobile phase using the same columns and detector as described above. The area under the curve of the polymer peak was used to calculate the polymer concentration after calibration with a PBD-*b*-PEO standard curve.

The concentration of dye in the fluorescent polymersome dispersion was determined by fluorescence spectroscopy. The dispersion was diluted in *N,N*-dimethylformamide (final polymersome dispersion volume fraction 5%, v/v). The samples were subsequently centrifuged at 5000 *x g* for 2 min. The fluorescence emission of the supernatant was determined at 510 nm (excitation wavelength 413 nm). The HPTS concentration in the polymersome dispersion was calculated with a HPTS standard curve in the same phosphate buffer/*N,N*-dimethylformamide mixture.

#### 4.3.2 Quantification of residual dichloromethane

The dichloromethane content was determined using headspace gas chromatography coupled to a flame ionization detector. The samples were mounted by headspace injection (TurboMatrix HS 40, Perkin Elmer, Waltham, MA) and measured by HP6890 (Hewlett Packard, Palo Alto, CA) equipped with a TR-FFAP column (50 m × 0.32 mm × 0.5 μm, Thermo Fisher Scientific). The samples were heated for 20 min at 35°C and the headspace needle and transfer line were set to 90 and 120°C, respectively. Samples were injected for 0.04 min at a total flow of 40.3 mL/min using a split flow of 36.5 mL/min (split ratio 20:1, pressure 90 kPa). Helium (Pangas, Dagmarsellen, Switzerland) was selected as the carrier gas. The pressure in the column was constant (90 kPa) and the flow 1.8 mL/min. The oven temperature was set to 45°C for the first 2 min and gradually increased to 130°C over the next 6 min. The FID detector was heated to 250°C with a hydrogen flow of 55 mL/min and an airflow of 300 mL/min.

#### 4.3.3 Size and morphology

The diameters of the PS-*b*-PEO (PS fragment size range: 2800 – 4100) and PBD-*b*-PEO polymersomes, liposomes, and microparticles were determined using the Mastersizer2000 laser diffraction particle size analyzer (Malvern Instruments, Malvern, UK). The results are presented as volume distribution (Supplementary Table S1, Supplementary Figure S14). The morphology of the PS-*b*-PEO polymersomes was further analyzed by cryo-scanning electron microscopy (SEM). For cryo-SEM, polymersome dispersion in citric acid solution 250 mM at pH 2.0 was loaded on a 6-mm aluminum carrier (sample thickness 200 µm). In a high-pressure freezer (HPM100, Bal-Tec, Vienna, Austria), the sample was vitrified at −120°C and mechanically broken (BAF-060, Bal-Tec). The resulting surface was coated with tungsten, transferred using a VCT100 (Bal-Tec), and imaged by SEM (Leo-1530, Zeiss, Oberkochen, Germany) at −120°C in accordance with the literature.^[56]^

### 4.4 Liposome preparation and characterization

Liposomes composed of (i) 1,2-dipalmitoyl-*sn*-glycero-3-phosphocholine (DPPC, LIPOID, Ludwigshafen, Germany) or 1,2-distearoyl-*sn*-glycero-3-phosphocholine (DSPC, LIPOID), (ii) cholesterol (Sigma Aldrich Chemie) and (iii) 1,2-distearoyl-*sn*-glycero-3-phosphoethanol-amine-*N*-[methoxy(PEO)-2000] (DSPE-PEO, LIPOID) at 54:45:1 mol% were prepared by the film hydration method.

For the DSPC liposomes, 8.2 mg cholesterol, 20.2 mg DSPC, and 1.3 mg DSPE-PEO were added as chloroform stock solutions to a glass vial. For the DPPC liposomes, 19.7 mg DPPC, 8.7 mg cholesterol, and 1.4 mg DSPE-PEO, were added as chloroform stock solutions to a glass vial. The organic solvent was subsequently removed by nitrogen flow for 2 h and storage in a vacuum desiccator overnight. The dried film was hydrated with 1 mL of citric acid solution 250 mM at pH 2.0 at 300 mOsmol/kg (lipid concentration = 29.8 mg/mL) while heating to 55°C and slowly mixing until the lipid film was not visible anymore. The size was determined as laid out in section 4.3.3.

### 4.5 *In vitro* ammonia uptake studies

#### 4.5.1 Ammonia uptake by polymersomes and liposomes

The transmembrane pH-gradient was generated in side-by-side diffusion cells (PermeGear Inc., Hellertown, PA) by dilution in alkaline phosphate buffer (final phosphate concentration 50 mM, pH 6.8, osmolality 300 mOsmol/kg) in the absence and presence of the bile salts sodium cholate (SC), sodium deoxycholate (SDC; bile salts for microbiology, Sigma Aldrich Chemie, 1:1 (m/m) mixture of SC and SDC), and sodium taurocholate (STC, abcr, Zug, Switzerland) at 25/25/0 mM (liposome, PBD-b-PEO, and hypoosmolal PS-*b*-PEO experiments), 12.5/12.5/0 (digestive enzyme experiment), or 30/30/30 mM (other experiments). In the hypoosmolal conditions, the osmolality was set to 160 mOsmol/kg by modifying the phosphate concentration to 10 mM and adjusting the sodium chloride concentration accordingly. In the hyperosmolal conditions, the osmolality was set to 620 mOsmol/kg by adjusting the sodium chloride concentration. In the digestive enzyme experiment, trypsin from porcine pancreas 1 mg/mL (approximately 10 000 IU/mL, Sigma Aldrich Chemie), α-chymotrypsin from bovine pancreas (Type II) 1 mg/mL (approximately 40 IU/mL, Sigma Aldrich Chemie), lipase from porcine pancreas (Type II) 3 mg/mL (approximately 300 IU/mL, Sigma Aldrich Chemie) were added to the bile salt-containing phosphate buffer. In the high potassium experiment, potassium chloride 200 mM was added to the potassium phosphate buffer 50 mM (final potassium concentration 250 mM).

The dual-chamber system was separated by a track etch polycarbonate membrane (pore size = 100 nm, Sterlitech, Kent, WA), physically isolating the vesicles on one side. At 37°C, the lipid or polymer and the ammonia concentrations within the diffusion cells were 1.75 mg/mL and 1.5 mM, respectively. At the allotted time points, aliquots were taken from the vesicle-free compartment. The ammonia concentration of the samples was determined by the Berthelot reaction.^[13]^ The ammonia uptake was quantified using equation 1 with the total mass of sequestrant being the total mass of lipid or polymer:

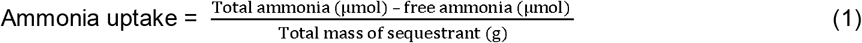

#### 4.5.2 Ammonia uptake with microparticles

To evaluate their ammonia uptake, zeolite (Sigma Aldrich Chemie), poly(methacrylic acid) (PMAA, Lewatit CNP-105, Sigma Aldrich Chemie), mesostructured silica (Sigma Aldrich Chemie), or activated charcoal (Norit, Cabot, Alpharetta, GA) microparticles were incubated at 30 mg/mL in phosphate buffer 50 mM (pH 6.8, 37°C) with an ammonia concentration of 1.5 mM under orbital shaking. At the allotted time points, the dispersions were filtered with a 0.2 µm syringe filter and the ammonia concentration of the samples was determined by the Berthelot reaction. The ammonia uptake was quantified using equation 1 with the total mass of sequestrant being the total mass of microparticles.

The cation competition experiments with zeolite microparticles (20 mg/mL) were conducted similarly with a modified buffer (HEPES 100 mM at pH 7.0 supplemented with 50 to 2000 mM sodium, potassium, or calcium) and an incubation time of 30 min at 37°C.

### 4.6 *In vitro* stability experiments in dietary fiber hydrogels

PS-*b*-PEO(2000) polymersomes with PS blocks between 3300 and 4500 were used for the dietary hydrogel experiments.

PS-*b*-PEO polymersomes were incubated in an isotonic phosphate buffer 10 mM at pH 6.8 with different Metamucil^®^ (Metamucil^®^ Regular, Procter & Gamble Switzerland, Petit-Lancy, Switzerland) concentrations (5-50%, m/m) for 24 h at 37°C. Subsequently, the hydrogel was diluted approx. 1:19 (m/v) in isotonic phosphate buffer 50 mM at pH 6.8 (final PS-*b*-PEO polymer concentration: 1 mg/mL; final HPTS concentration for the HTPS-containing transmembrane pH gradient polymersomes and the HTPS/DPX-containing polymersomes: 50 µM and 30 µM, respectively) containing 1.5 mM ammonia (non-fluorescent polymersomes) or no ammonia (fluorescent polymersomes). After 3 h incubation at 37°C under orbital shaking, the dispersions were centrifuged for 5 min at 4500 *x g*. The samples were filtered with a 0.2 µm syringe filter.

For the non-fluorescent polymersomes, the ammonia concentration of the filtered solution was determined using the L-glutamate dehydrogenase (GLDH)-based enzymatic ammonia kit (AM1015, Randox Laboratories, Crumlin, UK). For the fluorescent transmembrane pH-gradient polymersomes, the fluorescence emission intensities at 510 nm at an excitation wavelength of 455 nm (pH-dependent excitation wavelength) and 413 nm (pH-independent wavelength) were measured (Tecan Infinite 200 Pro). The fluorescence emission intensity ratio at these excitation wavelengths was subsequently calculated and compared with the non-pH-gradient control. For the DPX/HPTS-containing polymersomes, the fluorescence emission intensity at 510 nm at an excitation wavelength of 413 nm (pH-independent wavelength) was measured (Tecan Infinite 200 Pro). To account for incomplete extraction from the hydrogel matrix, the fluorescence intensity of DPX/HPTS-containing polymersomes were corrected with the extraction efficiency of HPTS-only polymersomes (Supplementary Figure S15).

### 4.7 Polymersome ammonia assay

PS-*b*-PEO(2000) polymersomes with PS blocks between 2800 and 4500 were used for the polymersome-based ammonia assay experiments.

The fluorescence excitation spectra of HPTS at 100 µM in isotonic citric acid solution 5.5 mM (pH 2.0-5.7) or isotonic phosphate buffer 50 mM (pH 6.8-11.0) was determined at a fixed emission wavelength of 510 nm (Tecan Infinite 200 Pro) at room temperature.

The samples, standards, or blank were co-incubated with polymersomes in isotonic phosphate buffer 50 mM at pH 7.4 at room temperature for 10 min (unless stated otherwise). The volume fraction of the sample, standard, or blank was 17% and 43% (v/v) per well with a final HPTS concentration of 16 and 40 µM, respectively. The fluorescence emission at 510 nm was measured at an isosbestic (413 nm) and at a pH-dependent excitation wavelength (455 nm) to determine the fluorescence emission ratio (*i.e.*, fluorescence emission at pH-dependent excitation wavelength normalized to the emission at the isosbestic excitation wavelength). The fluorescence emission ratio of the ammonia standards was corrected by subtracting the one of the blank (corrected fluorescence emission ratio). The accuracy was defined as the percentage of the mean of the measurements normalized to the nominal ammonia concentration. The precision was defined as coefficient of variation (*i.e.*, standard deviation normalized to mean). The lower limit of quantification (LOQ) was determined using equation 2 which relates to the current guidelines for bioanalytical method validation of the European Medicine’s Agency (EMA, EMEA/CHMP/EWP/192217/2009, legal effective date 2012/02/01)

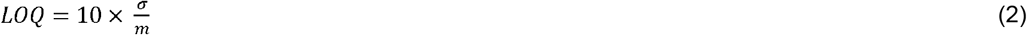

where σ stands for the standard deviation of the intercept and m for the slope of the standard curve.

#### 4.7.1 Selectivity

The ammonia concentration of a 100 µM ammonia solution (17%, v/v) was determined in the presence of a mixture of drugs and metabolites (dopamine (TCI), levofloxacin (Sigma Aldrich Chemie), propranolol (Sigma Aldrich Chemie), glycine (Acros), alanine (Roth, Arlesheim, Switzerland), each at 500 µM), or di- (200 µM, Sigma Aldrich Chemie) or trimethylamine (100 µM, TCI) (final volume fraction 17% (v/v) of the solution containing the potentially interfering substance). The ammonia concentration was quantified as described above by comparison to an ammonia standard curve in isotonic phosphate buffer 50 mM at pH 7.4.

#### 4.7.2 Time dependence

Time dependence of ammonia sensing with HPTS-containing PS-*b*-PEO polymersomes. Normalized fluorescence emission ratio of fluorescent polymersomes were exposed to ammonium chloride 50 or 200 µM in both the low and high standard volume fraction set-ups (17 or 43% (v/v) with HPTS concentration of 16 and 40 µM) for 5, 10, or 30 min (separate wells for each time point) before the fluorescence emission intensity was measured as described above. The fluorescence emission ratio at 10 or 30 min was normalized to the one at 5 min.

#### 4.7.3 Stability at 4°C

Fluorescent polymersomes were purified with phosphate buffer 50 mM at pH 7.4 as described above and stored for 5 months at 4°C protected from light. The polymersome assay was conducted as described above and the fluorescence emission ratio of a phosphate buffer blank was subtracted (corrected fluorescence emission ratio).

#### 4.7.4 Phenylalanine sensing

Commercial phenylalanine ammonia lyase from *Rhodotorula glutinis* (EC number 4.3.1.5, grade I, activity: 0.8-2.0 units/mg protein (1 unit converts 0.001 mmol phenylalanine per min at pH 8.5 at 30°C), Sigma Aldrich Chemie), was purified using centrifugal filtration (cut-off 30 kDa, Vivaspin 500, Sartorius AG, Goettingen, Germany) eight times for 2 min at 15’000 × g at 4°C. Phenylalanine ammonia lyase (0.026 mg/mL) was incubated with different phenylalanine solutions (0 - 1.5 mM, volume fraction 50%) in tris(hydroxymethyl)aminomethane 5 mM at pH 8.5 at 300 mOsmol/kg for 15 min at 30°C. The samples were subsequently incubated with fluorescent polymersomes as described above (volume fraction of sample 43%) and the fluorescence emission ratio measured as described above.

#### 4.7.5 Quantification in other body fluids

##### 4.7.5.1 Saliva, semen, urine, sweat

Saliva, semen, urine, sweat (all from healthy human donors, Lee BioSolutions, Maryland Heights, MO) were incubated with fluorescent polymersomes at final sample volume fractions of 0.17% (semen, urine) or 1.7% (saliva, sweat) for 10 min at room temperature (HPTS concentration 16 µM). The concentrations were calculated with an ammonia standard curve in isotonic phosphate buffer at pH 7.4. The samples were volumetrically diluted in isotonic phosphate buffer at pH 7.4 1/100 (semen, urine) or 1/10 (saliva, sweat) and measured with the GLDH assay.

##### 4.7.5.2 Intestinal fluid

Caecum fluid from healthy female Sprague Dawley rats (8-10 weeks old) was centrifuged for 10 min at 15’000 × *g* at 4°C. The supernatant was spiked with ammonium chloride standards in water (final volume fraction of standard 10%). The spiked caecum samples were incubated with fluorescent polymersomes at a final sample volume fraction of 0.5% for 10 min at room temperature. The fluorescence emission ratio was subsequently measured as described above, and the fluorescence emission ratio emission of a phosphate buffer blank was subtracted (corrected fluorescence emission ratio).

### 4.8. *In vivo* experiments with BDL rats

The animal experiments adhered to national guidelines and were approved by the local ethics committee of the Centre de Recherche du CHUM (CIPA, Montreal, QC, Canada). Secondary biliary cirrhosis was induced in male Sprague-Dawley rats (ca. 220 g) (Charles River Laboratories, St. Constant, QC, Canada) by surgical bile duct ligation (BDL). In brief, laparotomy was performed and the common bile duct was located, ligated, and resected using a dissection microscope as described previously.^[57]^

#### 4.8.1 Oral administration of polymersomes

In the first part of the oral study, the laxative effects of poly(ethylene glycol) (3350) (PEG, 1 g/kg, n = 3, Pegalax^®^, Aralez Pharmaceuticals, Mississauga, ON, Canada), sodium picosulfate (25 mg/kg, n = 19, *i.e.*, same animals used in the next paragraph, Dulcolax^®^ Picosulfate, Sanofi Aventis, Vernier, Switzerland), and the negative control (water, n = 6) were determined. The laxatives were diluted in water and gavaged once daily at 8.30 a.m. (5 mL/kg) in BDL rats at three weeks after surgery. Fresh stool samples were collected after 8 and 24 h and dried at 70°C overnight. The water loss of the stool samples after drying was quantified gravimetrically.

In the second part, the effects of an orally applied polymersome dispersion (1 g polymer/kg per day, diblock copolymers from Supplementary Table S7, individual polymersome batches were pooled and gavaged as is, *i.e.*, dispersed in citric acid solution 250 mM at pH 2.0) was assessed in BDL rats supplemented with sodium picosulfate (25 mg/kg per day). Three weeks after surgery, blood was sampled from the saphenous vein into heparin-containing tubes (Sandoz Canada, Boucherville, QC, final concentration approx. 20 IU/mL) and centrifuged at 1500 *x g* at 4°C for 7 min. The ammonia levels in the heparinized plasma were immediately measured using the PocketChem BA PA-4140 (20 µL sample, mode F6; PocketChem BA PA-4140, Arkray, Kyoto, Japan). Two groups with similar mean ammonia levels were created (n = 18-19 per group) which received sodium picosulfate (once daily 25 mg/kg) and polymersome dispersion in citric acid solution (twice daily 0.5 g polymer/kg) or citric acid solution alone (twice daily 0.24 g citric acid/kg, *i.e.*, corresponding to citric acid dose received by the polymersome group) by gavage twice daily (10.30 a.m. and 5 p.m.) for one week (total gavage volume: 15 mL/kg per day). A group of age-matched healthy rats (n = 6) served as control and received citric acid solution (twice daily 0.24 g/kg) by gavage for one week (15 mL/kg per day). At day seven, the second dose of polymersome dispersion or citric acid solution was administered at 1 p.m. and the rats were sacrificed at 3.30 p.m. Arterial blood was collected by cardiac puncture in heparin-containing tubes (final concentration 10 IU/mL) and centrifuged at 1500 *x g* at 4°C for 7 min. The ammonia levels in the heparinized plasma were immediately measured with the PocketChem BA as described above. Three and one deaths occurred in the polymersome and the citric acid group, respectively. All deaths occurred right after the gavage procedure.

#### 4.8.2 Quantification of ammonia in rat plasma

The polymersome assay was evaluated in rat plasma. BDL rats at four weeks after surgery and weight-matched healthy male Sprague-Dawley rats were used. Approximately 5 mL of arterial blood were collected by cardiac puncture in heparin-containing tubes (final concentration 10 IU/mL) and centrifuged at 1500 *x g* at 4°C for 7 min. The samples were frozen and stored at −80°C. After thawing on ice, the ammonia concentration was analyzed with the polymersome assay (compared with a standard curve in phosphate buffer at pH 7.4) as described above using a different plate reader (Tecan Spark^®^ 20M, Tecan), the enzymatic GLDH-based ammonia assay, and the PocketChem BA. Selected biochemical laboratory parameters (alanine transaminase, ALT, aspartate transaminase, AST, albumin, total bilirubin, alkaline phosphatase, AP, gamma-glutamyl transpeptidase, GGT) were analyzed from BDL plasma samples at the CHUM (Montréal, QC, Canada).

### 4.9 Commercial ammonia assays

In the GLDH enzymatic ammonia kit, the samples were measured based on the manufacturer’s instructions with a modified sample (20 µL), reagent buffer (200 µL), and the enzyme solution volume (2 µL) in order to adapt the protocol to a 96-well plate set-up.^[14,15]^ In brief, the ammonia-containing solution, blank, or calibrator was added to the reagent buffer (trometamol 150 mM at pH 8.6) in the well and incubated for 5 min at room temperature. After measuring the baseline absorbance at 340 nm (Tecan Infinite 200 Pro), the enzyme was added and the absorbance at the same wavelength was measured after 5 min incubation at room temperature. Based on an ammonium chloride standard curve (25-200 µM), the lowest concentration with acceptable accuracy (between 80-120%) and precision (CV below 20%) was determined (50 µM). The measurements below this value were set as 25 µM in accordance with the literature.^[14]^

To measure ammonia levels with the PocketChem BA, 20 µL of sample was applied onto the strip (Ammonia Test Kit II, Arkray) and measured in mode F6 according to the manufacturer’s instructions. To establish the ammonia standard curve, the ammonia standard from the Randox enzymatic ammonia assay (298 µM) was taken and diluted in water. The standards were measured in mode F6 according to the manufacturer’s instructions. In the TMA measurements with the PocketChem BA, TMA standards in phosphate buffer at pH 8.0 were prepared, applied on the strip, and measured after 1 min incubation.

In the ammonia quantification using the Berthelot reaction, the ammonia-containing sample was added to equivolumetric amounts of alkaline sodium hypochlorite and phenol nitroprusside solution (both from Sigma Aldrich Chemie). After 25 min of incubation at room temperature, the absorbance was measured at 636 nm (Tecan Infinite 200 Pro).

### 4.10 Statistical analysis

The statistical calculations were carried out by SigmaPlot (version 13.0) and Microsoft Excel 2016 (determination of R^2^). Three or more groups were compared using one-way ANOVA (Holm-Sidak test) assuming a normal distribution of the data. In ammonia uptake kinetics with side-by-side diffusion cells, the time point at 24 h was compared between groups with this test. Two groups were compared using an unpaired t-test. A p-value of <0.05 was deemed statistically significant.

## Acknowledgements

S.M. gratefully acknowledges a doctoral scholarship from the Swiss Chemical Industry (SSCI). We further acknowledge generous funding from the Swiss National Science Foundation (2-77082-16). We further thank Julia Müller, Dr. Valentina Agostoni, Dr. Meriam Kabbaj, Dr. Vincent Forster, Prof. Dr. Davide Brambilla, Dr. Jong-Ah Kim, Nevena Paunovic, Prof. Dr. Georgios Sotiriou, Dr. Anastasia Spyrogianni, Dr. Charlotte Gourmel, and Olha Wuerthinger for their support. Our thanks also go to Prof. Dr. Peter Walde for insightful discussions. Finally, we thank Susanne Freedrich (EPIC) for providing caecum content and scopeM for the assistance in the cryo-SEM experiment.

## References

[1] E. F. M. Wijdicks, N. Engl. J. Med. 2016, 375, 1660.

[2] S. Matoori, J.-C. Leroux, Adv. Drug Deliv. Rev. 2015, 90, 55.

[3] H. Vilstrup, P. Amodio, J. Bajaj, J. Cordoba, P. Ferenci, K. D. Mullen, K. Weissenborn, P. Wong, Hepatology 2014, 60, 715.

[4] J. P. Ong, A. Aggarwal, D. Krieger, K. A. Easley, M. T. Karafa, F. Van Lente, A. C. Arroliga, K. D. Mullen, Am. J. Med. 2003, 114, 188.

[5] J. Häberle, N. Boddaert, A. Burlina, A. Chakrapani, M. Dixon, M. Huemer, D. Karall, D. Martinelli, P. S. Crespo, R. Santer, A. Servais, V. Valayannopoulos, M. Lindner, V. Rubio, C. Dionisi-Vici, Orphanet J. Rare Dis. 2012, 7, 32.

[6] G. A. Diaz, L. S. Krivitzky, M. Mokhtarani, W. Rhead, J. Bartley, A. Feigenbaum, N. Longo, W. Berquist, S. A. Berry, R. Gallagher, U. Lichter-Konecki, D. Bartholomew, C. O. Harding, S. Cederbaum, S. E. McCandless, W. Smith, G. Vockley, S. A. Bart, M. S. Korson, D. Kronn, R. Zori, J. L. Merritt, S. C S Nagamani, J. Mauney, C. Lemons, K. Dickinson, T. L. Moors, D. F. Coakley, B. F. Scharschmidt, B. Lee, Hepatology 2013, 57, 2171.

[7] V. R. Patwardhan, Z. G. Jiang, Y. Risech-Neiman, G. Piatkowski, N. H. Afdhal, K. Mukamal, M. P. Curry, E. B. Tapper, J. Clin. Gastroenterol. 2016. 5, 345.

[8] J. M. Vierling, M. Mokhtarani, R. S. Brown, P. Mantry, D. C. Rockey, M. Ghabril, R. Rowell, M. Jurek, D. F. Coakley, B. F. Scharschmidt, Clin. Gastroenterol. Hepatol. 2016, 14, 903.

[9] C. Bachmann, Eur. J. Pediatr. 2003, 162, 410.

[10] R. Posset, A. Garcia-Cazorla, V. Valayannopoulos, E. L. Teles, C. Dionisi-Vici, A. Brassier, A. B. Burlina, P. Burgard, E. Cortès-Saladelafont, D. Dobbelaere, M. L. Couce, J. Sykut-Cegielska, J. Häberle, A. M. Lund, A. Chakrapani, M. Schiff, J. H. Walter, J. Zeman, R. Vara, S. Kölker, A. individual contributors of the E.-I. consortium, J. Inherit. Metab. Dis. 2016, 3, 661.

[11] I. Seiden-Long, K. Schnabl, W. Skoropadyk, N. Lennon, A. McKeage, Clin. Biochem. 2014, 47, 1116.

[12] R. Goggs, S. Serrano, B. Szladovits, I. Keir, R. Ong, D. Hughes, Vet. Clin. Pathol. 2008, 37, 198.

[13] V. Forster, R. D. Signorell, M. Roveri, J.-C. Leroux, Sci. Transl. Med. 2014, 6, 258ra141.

[14] V. Agostoni, S. H. Lee, V. Forster, M. Kabbaj, C. R. Bosoi, M. Tremblay, M. Zadory, C. F. Rose, J.-C. Leroux, Adv. Funct. Mater. 2016, 26, 8382.

[15] G. Giacalone, S. Matoori, V. Agostoni, V. Forster, M. Kabbaj, S. Eggenschwiler, M. Lussi, A. De Gottardi, N. Zamboni, J.-C. Leroux, J. Control. Release 2018, 27, 57.

[16] E. Moroz, S. Matoori, J.-C. Leroux, Adv. Drug Deliv. Rev. 2016, 101, 108.

[17] T. T. Kararli, Biopharm. Drug Dispos. 1995, 16, 351.

[18] K. T. Kim, J. J. L. M. Cornelissen, R. J. M. Nolte, J. C. M. van Hest, Adv. Mater. 2009, 21, 2787.

[19] R. S. M. Rikken, H. Engelkamp, R. J. M. Nolte, J. C. Maan, J. C. M. van Hest, D. A. Wilson, P. C. M. Christianen, Nat. Commun. 2016, 7, 12606.

[20] J.-F. Le Meins, O. Sandre, S. Lecommandoux, Eur. Phys. J. E. Soft Matter 2011, 34, 1.

[21] H. Bermudez, A. K. Brannan, D. A. Hammer, F. S. Bates, D. E. Discher, Macromolecules 2002, 35, 8203.

[22] D. Richter, R. Zorn, B. Farago, B. Frick, L. J. Fetters, Phys. Rev. Lett. 1992, 68, 71.

[23] T. G. Fox, P. J. Flory, J. Polym. Sci. 1954, 14, 315.

[24] C. R. Bosoi, C. Parent-Robitaille, K. Anderson, M. Tremblay, C. F. Rose, Hepatology 2011, 53, 1995.

[25] R. Schubert, H. Jaroni, J. Schoelmerich, K. H. Schmidt, Digestion 1983, 28, 181.

[26] K. Andrieux, L. Forte, S. Lesieur, M. Paternostre, M. Ollivon, C. Grabielle-Madelmont, Eur. J. Pharm. Biopharm. 2009, 71, 346.

[27] C. J. O’Connor, R. G. Wallace, K. Iwamoto, T. Taguchi, J. Sunamoto, Biochim. Biophys. Acta - Biomembr. 1985, 817, 95.

[28] R. N. Rowland, J. F. Woodley, Biochim. Biophys. Acta - Lipids Lipid Metab. 1980, 620, 400.

[29] O. Zumbuehl, H. G. Weder, Biochim. Biophys. Acta - Biomembr. 1981, 640, 252.

[30] M. Kokkona, P. Kallinteri, D. Fatouros, S. G. Antimisiaris, Eur. J. Pharm. Sci. 2000, 9, 245.

[31] V. Pata, F. Ahmed, D. Discher, N. Dan, Langmuir 2004, 20, 3888.

[32] M. A. Jama, H. Yücel, Sep. Sci. Technol. 1989, 24, 1393.

[33] M. R. Weir, G. L. Bakris, D. A. Bushinsky, M. R. Mayo, D. Garza, Y. Stasiv, J. Wittes, H. Christ-Schmidt, L. Berman, B. Pitt, N. Engl. J. Med. 2015, 37, 211.

[34] G. D. Zuidema, D. Cullen, R. S. Kowalczyk, E. F. Wolfman, Arch. Surg. 1963, 87, 296.

[35] M. P. de la C. Moreno, M. Oth, S. Deferme, F. Lammert, J. Tack, J. Dressman, P. Augustijns, J. Pharm. Pharmacol. 2006, 5, 1079.

[36] L. Kalantzi, K. Goumas, V. Kalioras, B. Abrahamsson, J. B. Dressman, C. Reppas, Pharm. Res. 2006, 23, 165.

[37] D.-A. Hallbäck, M. Jodal, M. Mannischeff, O. Lundgren, Acta Physiol. Scand. 1991, 143, 271.

[38] Z. Song, Y. Huang, V. Prasad, R. Baumgartner, S. Zhang, K. Harris, J. S. Katz, J. Cheng, ACS Appl. Mater. Interfaces 2016, 8, 17033.

[39] L. Zhang, A. Eisenberg, Science 1995, 268, 1728.

[40] E. Ruel-Gariépy, G. Leclair, P. Hildgen, A. Gupta, J.-C. Leroux, J. Control. Release 2002, 8, 373.

[41] Ž. Pavelić, N. Škalko-Basnet, J. Filipović-Grčić, A. Martinac, I. Jalšenjak, J. Control. Release 2005, 10, 34.

[42] S. Mourtas, S. Fotopoulou, S. Duraj, V. Sfika, C. Tsakiroglou, S. G. Antimisiaris, Colloids Surfaces B Biointerfaces 2007, 55, 212.

[43] R. M. Straubinger, D. Papahadjopoulos, K. Hong, Biochemistry 1990, 29, 4929.

[44] P. Tiefenboeck, J. A. Kim, F. Trunk, T. Eicher, E. Russo, A. Teijeira, C. Halin, J.-C. Leroux, ACS Nano 2017, 11, 7758.

[45] Z. Horak, J. Kolarik, M. Sipek, V. Hynek, F. Vecerka, J. Appl. Polym. Sci. 1998, 6, 2615.

[46] X. Liang, G. Mao, K. Y. S. Ng, J. Colloid Interface Sci. 2004, 278, 53.

[47] M. P. Desai, V. Labhasetwar, E. Walter, R. J. Levy, G. L. Amidon, Pharm. Res. 1997, 14, 1568.

[48] J. Parmentier, M. M. M. Becker, U. Heintz, G. Fricker, Int. J. Pharm. 2011, 405, 210.

[49] S. Hu, M. Niu, F. Hu, Y. Lu, J. Qi, Z. Yin, W. Wu, Int. J. Pharm. 2013, 441, 693.

[50] T. Teerlink, M. W. T. Hennekes, C. Mulder, H. F. H. Brulez, J. Chromatogr. B Biomed. Sci. Appl. 1997, 69, 269.

[51] R. Obeid, H. M. Awwad, Y. Rabagny, S. Graeber, W. Herrmann, J. Geisel, Am. J. Clin. Nutr. 2016, 103, 703.

[52] A. C. Chakrabarti, I. Clark-Lewis, P. R. Harrigan, P. R. Cullis, Biophys. J. 1992, 61, 228.

[53] R. F. Chen, Arch. Biochem. Biophys. 1974, 160, 106.

[54] N. Blau, F. J. van Spronsen, H. L. Levy, Lancet 2010, 376, 1417.

[55] S. Hocine, D. Cui, M.-N. Rager, A. Di Cicco, J.-M. Liu, J. Wdzieczak-Bakala, A. Brûlet, M.-H. Li, Langmuir 2013, 29, 1356.

[56] L. Isa, F. Lucas, R. Wepf, E. Reimhult, Nat. Commun. 2011, 2, 438.

[57] C. R. Bosoi, M. M. Oliveira, R. Ochoa-Sanchez, M. Tremblay, G. A. Ten Have, N. E. Deutz, C. F. Rose, C. Bemeur, Metab. Brain Dis. 2017, 32, 513.

