## Supplementary Information for "An investigation of PS-*b*-PEO polymersomes for the oral treatment and diagnosis of hyperammonemia"

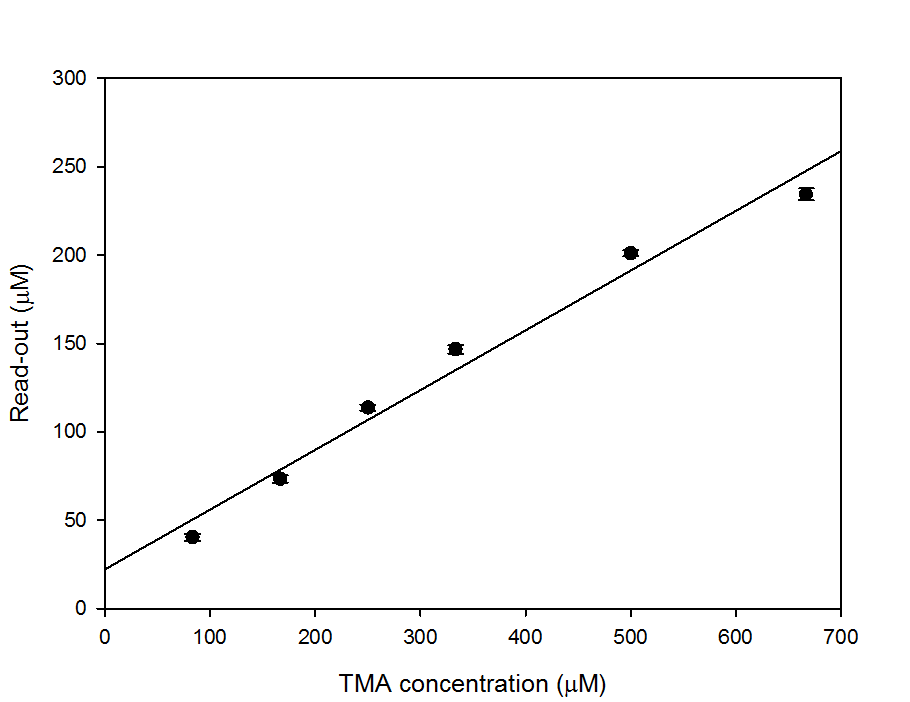

**Supplementary Figure S1.** TMA sensing with PocketChem BA. PocketChem BA read-out upon application of TMA-containing isotonic phosphate buffer at pH 8.0 onto strip. Incubation time 1 min; sample volume 20 µL. All results as mean ± SD (n = 3).

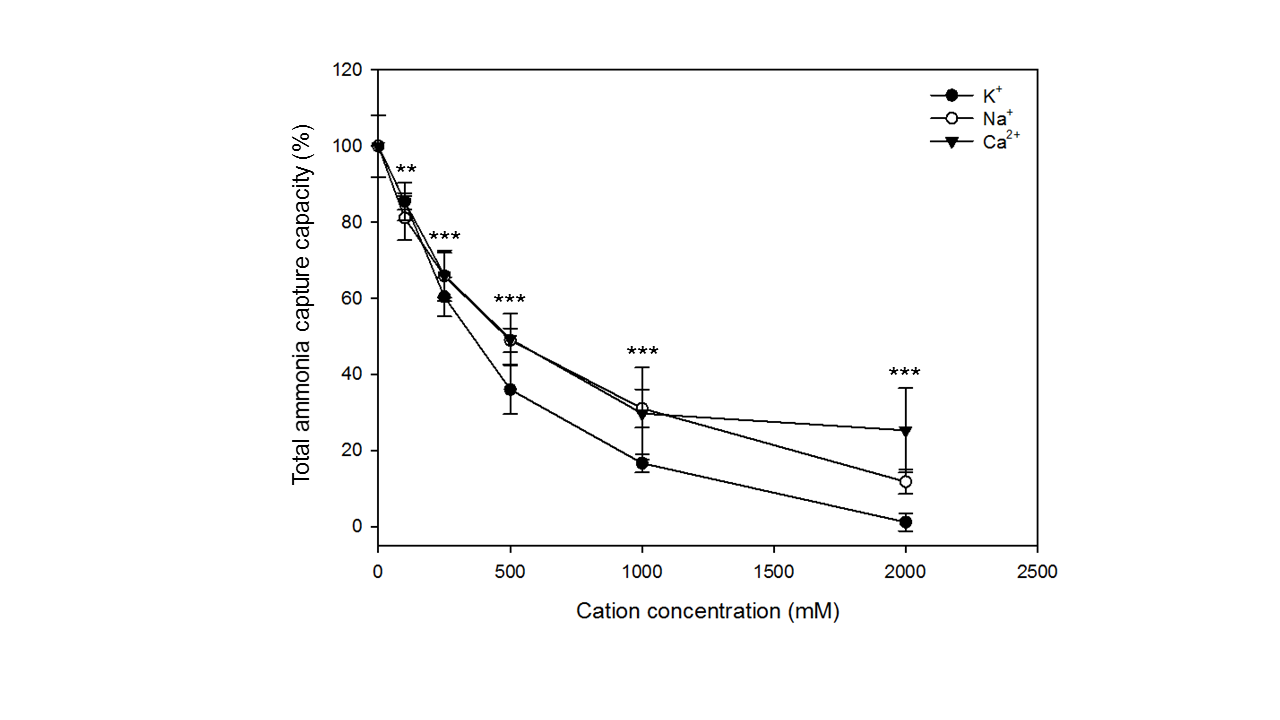

**Supplementary Figure S2.** Competitive binding of ammonia and cations with zeolites. Normalized ammonia uptake of zeolite microparticles at 20 mg/mL in the presence of increasing cation (potassium, sodium, calcium) concentrations. Buffer composition: HEPES buffer 100 mM at pH 7.0; ammonia concentration: 1.0 mM; n = 3-13. All results as means ± SD. **p < 0.01, and ***p < 0.001.

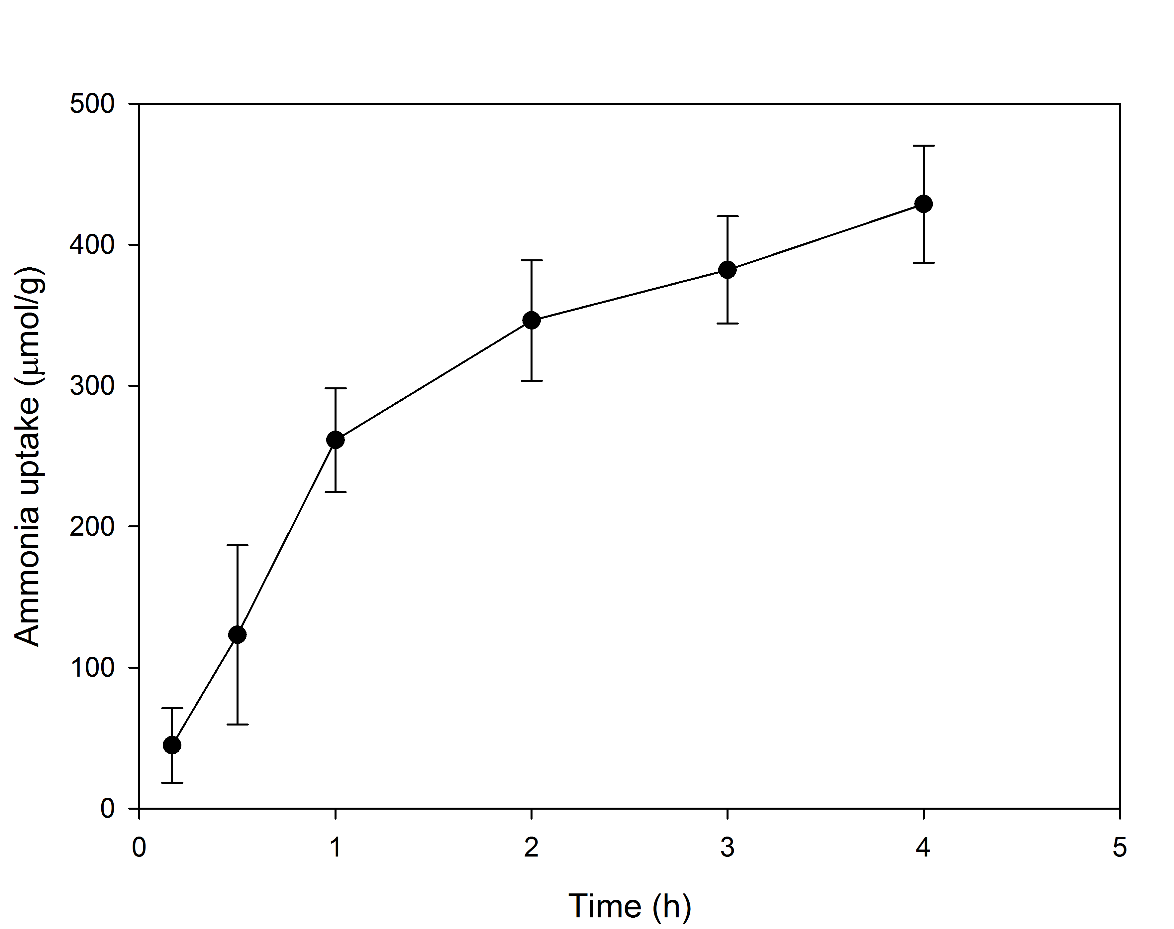

**Supplementary Figure S3.** Ammonia uptake in the presence of digestive enzymes. Ammonia uptake of PS(5000)-*b*-PEO(2000) corrected for ammoniagenic protein degradation upon incubation in a trypsin- (1.0 mg/mL), chymotrypsin- (1.0 mg/mL), lipase- (3.0 mg/mL), and bile salt-containing buffer at pH 6.8. Bile salt composition in isotonic phosphate buffer 50 mM at pH 6.8: cholate and deoxycholate, 12.5 mM each; polymer concentration: 1.75 mg/mL; ammonia concentration 1.5 mM; temperature 37°C. All results as means ± SD (n = 3).

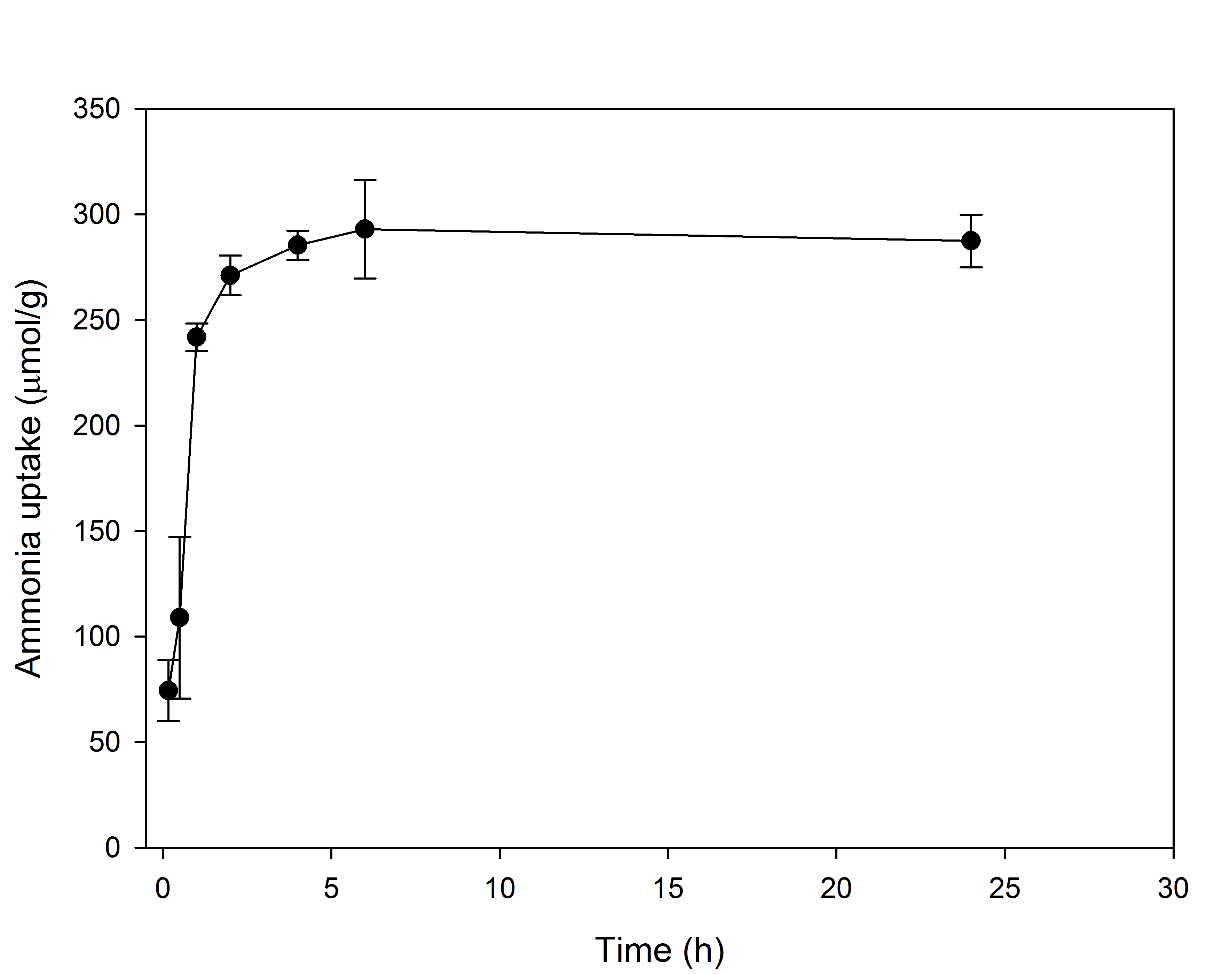

**Supplementary Figure S4.** Ammonia uptake in the presence of high potassium concentrations. Ammonia uptake of PS(2500)-*b*-PEO(2000) in a potassium chloride 250 mM and bile salt-containing phosphate buffer at pH 6.8. Bile salt composition in phosphate buffer 50 mM at pH 6.8: cholate, deoxycholate, and taurocholate, 30 mM each; polymer concentration: 1.75 mg/mL; ammonia concentration 1.5 mM; temperature 37°C. All results as means ± SD (n = 3).

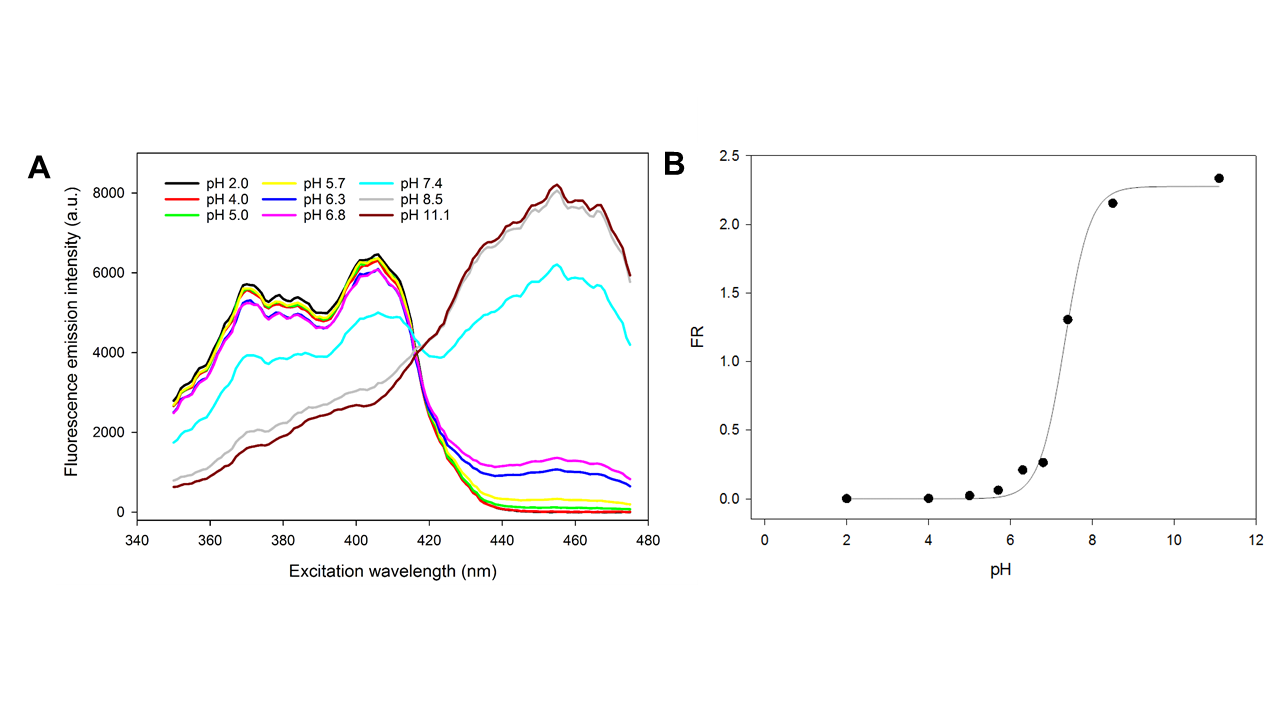

**Supplementary Figure S5.** Fluorescence emission of HPTS at different pH values. Fluorescence emission spectra (λ_em_ 510 nm) for free HPTS in solution (A). HPTS fluorescence emission ratio (FR) at different pH values in solution (B). FR refers to emission intensity at λ_em_ 510 nm for λ_ex_ 455 nm (pH-dependent λ_ex_) normalized to the emission at the same λ_em_ for λ_ex_ 413 nm (isosbestic λ_ex_). Buffer solutions: citrate buffer 5.5 mM (pH 2.0-5.7), phosphate buffer 50 mM (pH 6.3 – 7.4), or tris 5 mM (pH 8.5, 11.1), all at 300 mOsmol/kg. Dye concentration 100 µM.

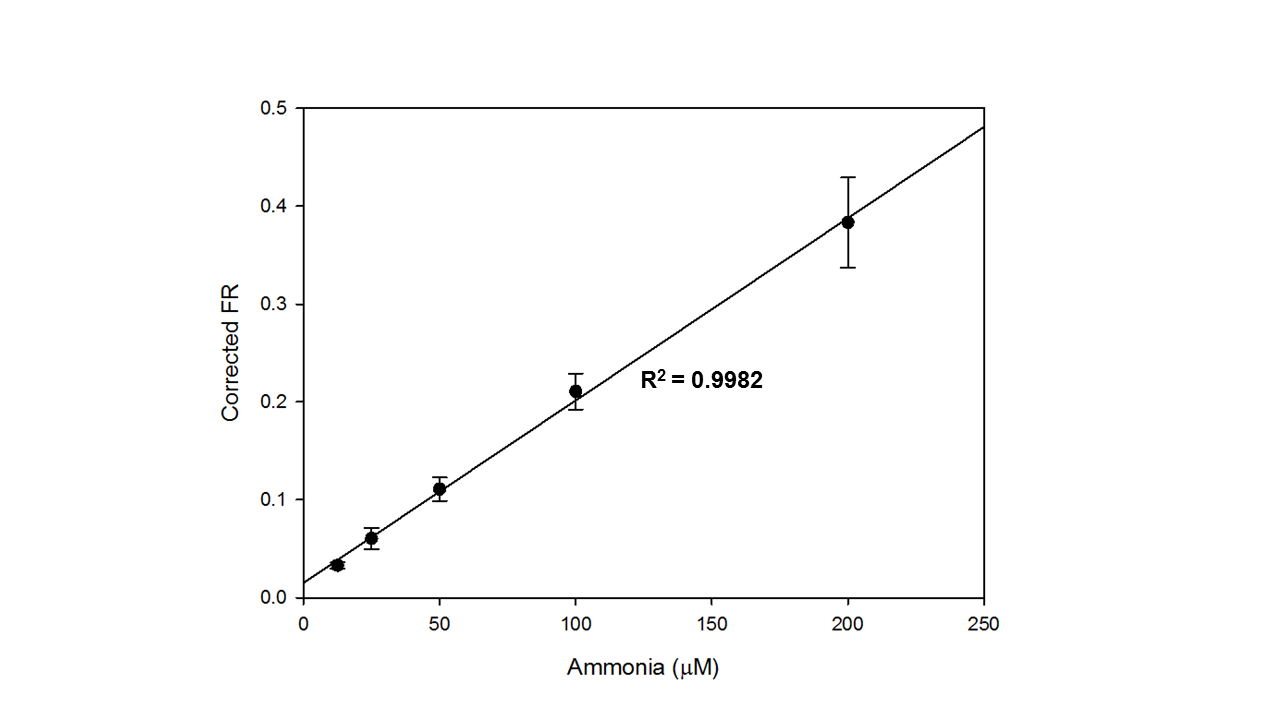

**Supplementary Figure S6.** Blank-corrected fluorescence emission ratio of HPTS-loaded polymersomes at different ammonia concentrations. HPTS concentration in final dispersion: 40 µM. Ammonia standard volume fraction: 43%. All results as mean ± SD (n = 3).

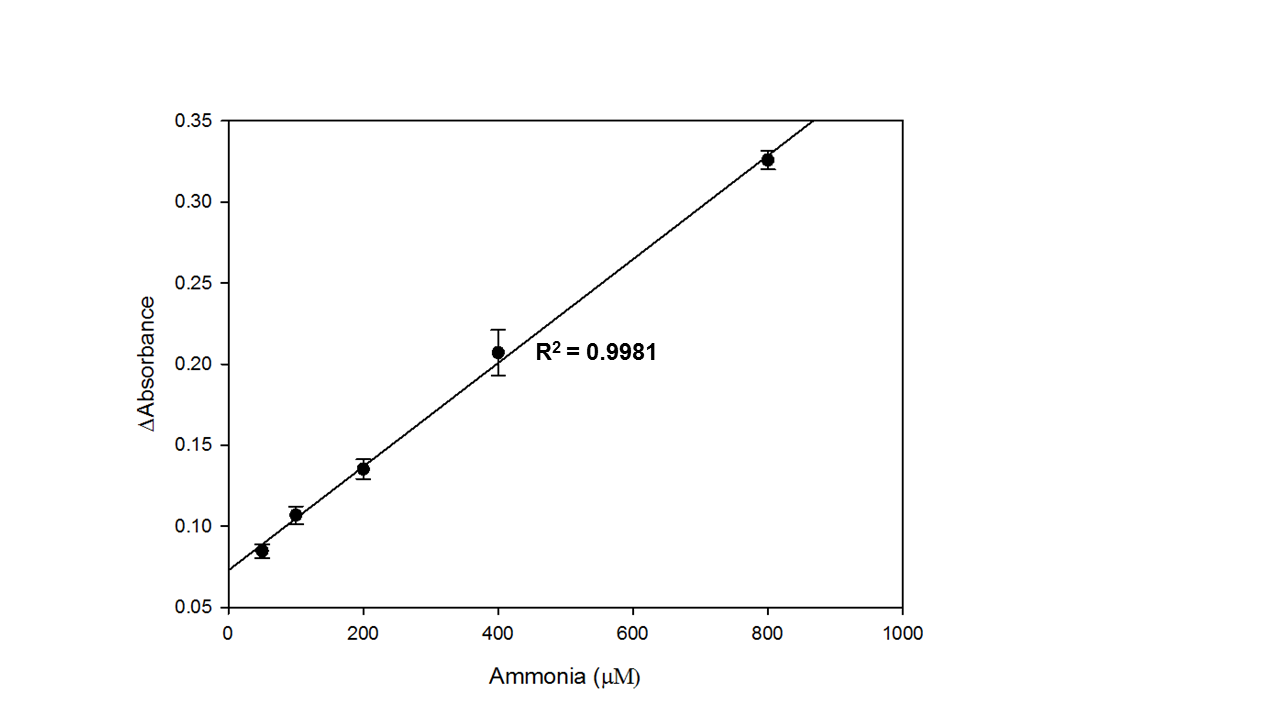

**Supplementary Figure S7.** Ammonia standard curve with GLDH-based ammonia assay. Baseline-corrected change in absorbance at 340 nm at different ammonia concentrations. GLDH catalyzes the conversion of alpha-ketoglutarate and ammonia to L-glutamate under the oxidation of a nicotinamide adenine dinucleotide phosphate (NADPH) molecule. The formation of the oxidized NADP^+^ was followed spectrophotometrically at 340 nm.[1] All results as mean ± SD (n = 8).

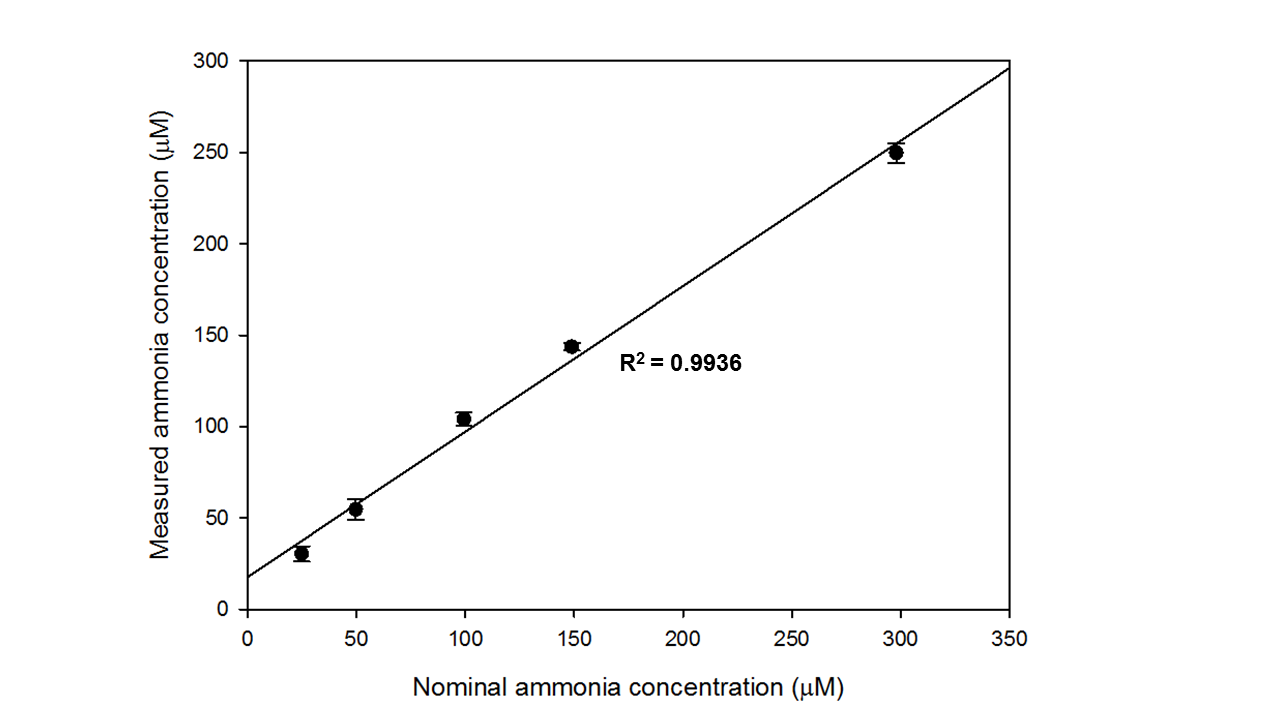

**Supplementary Figure S8.** Ammonia standard curve with PocketChem BA. PocketChem BA read-out of ammonia standards in water measured in mode F6. Incubation time 3 min; sample volume 20 uL. All results as mean ± SD (n = 3).

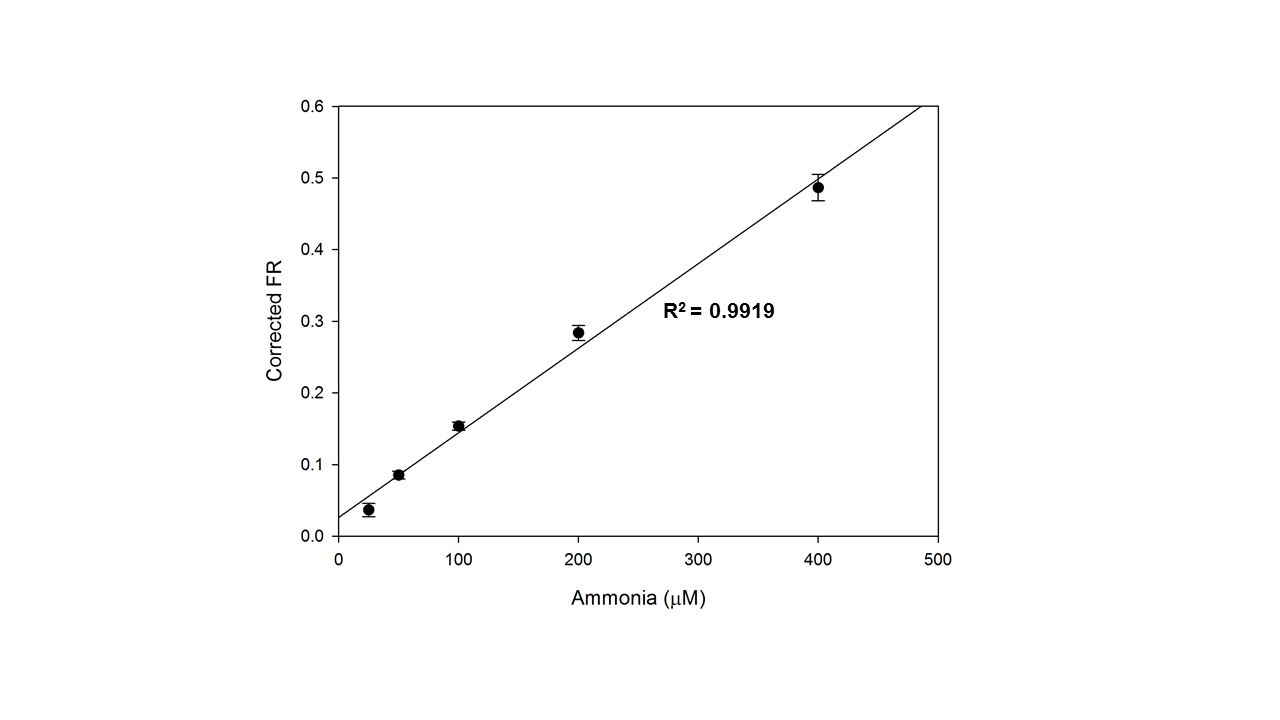

**Supplementary Figure S9.** Stability of HPTS-containing PS-*b*-PEO polymersomes stored at pH 7.4 and 4°C for five months. Blank-corrected fluorescence emission ratio at different ammonia concentrations (R^2^ = 0.9919). Inner phase: isotonic citrate buffer 5 mM at pH 5.5, outer phase: isotonic phosphate buffer 50 mM at pH 7.4. HPTS concentration in final dispersion: 16 µM. Volume fraction of ammonia standard: 17%. Corrected fluorescence emission ratio (FR): FR of blank subtracted from FR at given ammonia concentration. All results as mean ± SD (n = 3).

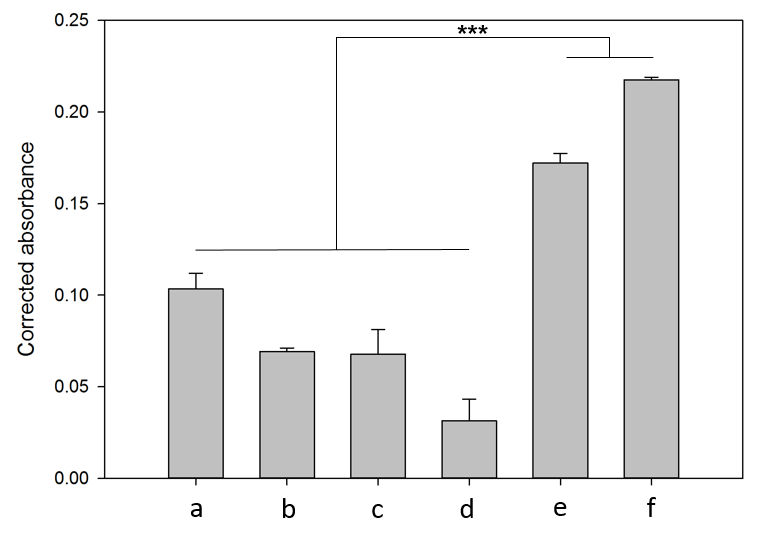

**Supplementary Figure S10.** Determination of hemolysis in healthy rat plasma. Corrected absorbance at 340 nm in the same six rats as in Figure 3C. All results as mean ± SD (n = 3-10). ***p < 0.001.

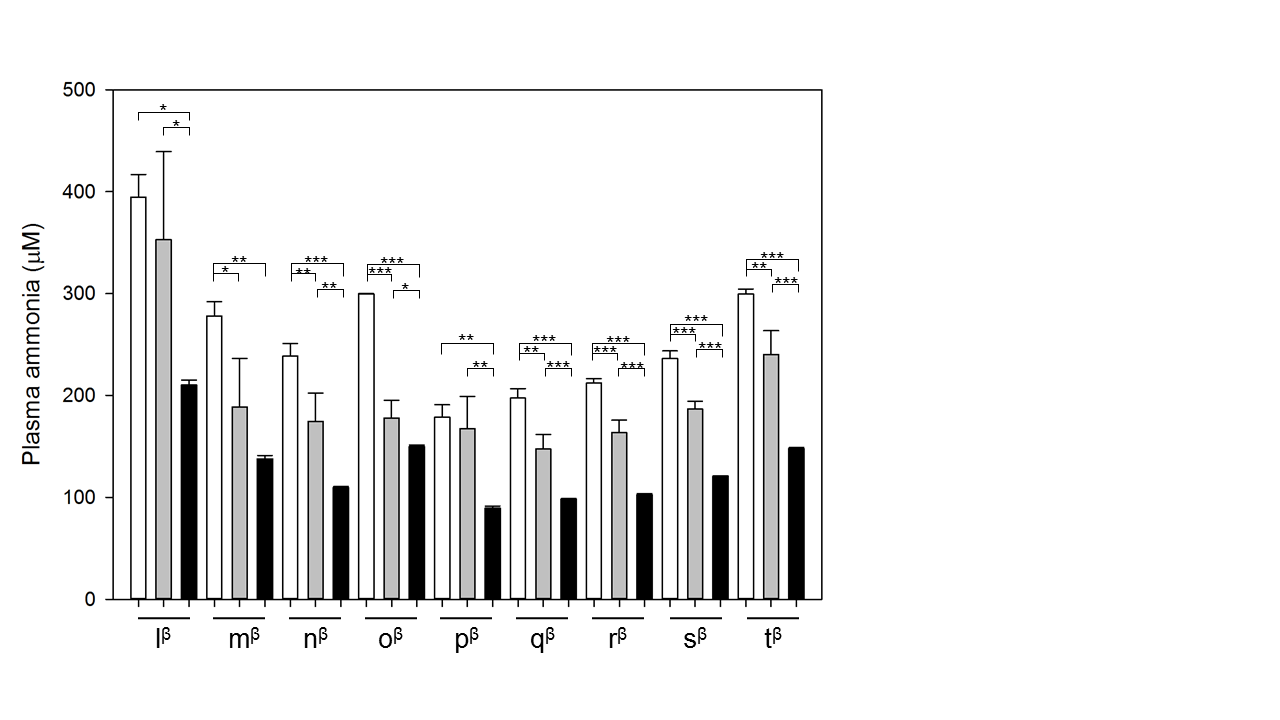

**Supplementary Figure S11.** Plasma ammonia quantification in BDL rats. Arterial plasma samples (cardiac puncture) from BDL (β, four weeks after surgery) male Sprague-Dawley rats measured with the polymersome assay (white), the GLDH-based ammonia assay by Randox (gray), and the PocketChem BA (black) (n = 3). Sample/standard volume fraction: 43%. HPTS concentration: 40 µM. Each letter represents one animal. All results as mean ± SD (n = 3). *p < 0.05, **p < 0.01, and ***p < 0.001.

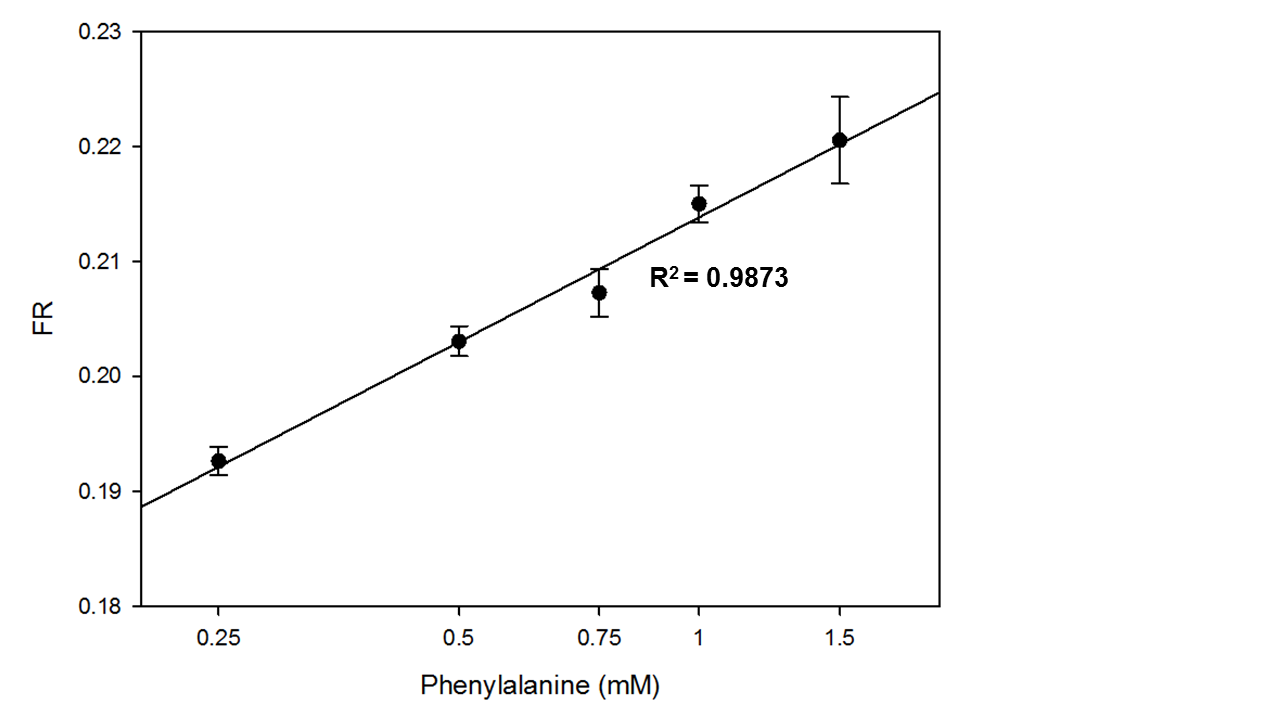

**Supplementary Figure S12.** L-phenylalanine ammonia lyase-mediated phenylalanine sensing with polymersome assay. Fluorescence emission ratio of HPTS-containing polymersomes with different L-phenylalanine solutions pre-exposed to L-phenylalanine ammonia lyase. Inner phase: isotonic citrate buffer 5 mM at pH 5.5, outer phase: isotonic phosphate buffer 50 mM at pH 7.4. L-phenylalanine solutions pre-exposed to L-phenylalanine ammonia lyase in isotonic tris buffer 5 mM at pH 8.5 for 15 min at 30°C. HPTS concentration in final dispersion: 40 µM. Sample volume fraction: 43%. All results as mean ± SD (n = 3).

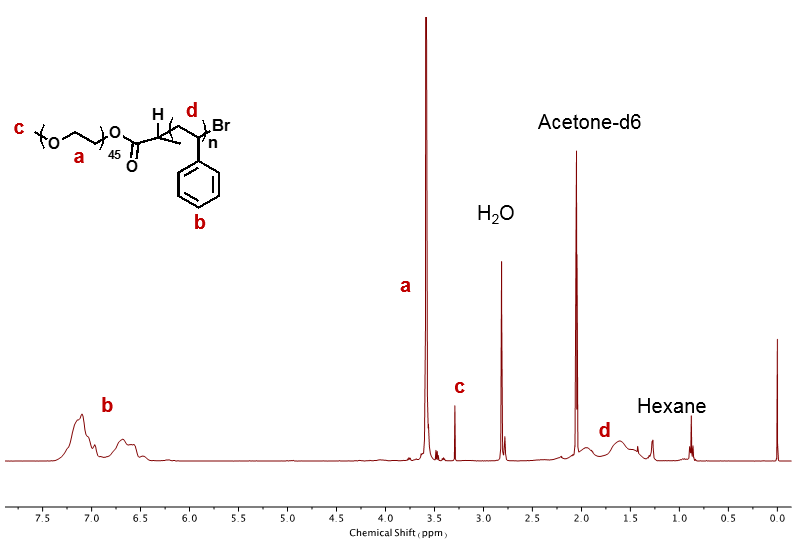

**Supplementary Figure S13.** ^1^H NMR spectrum of PS(4500)-*b*-PEO(2000) diblock copolymer in deuterated acetone.

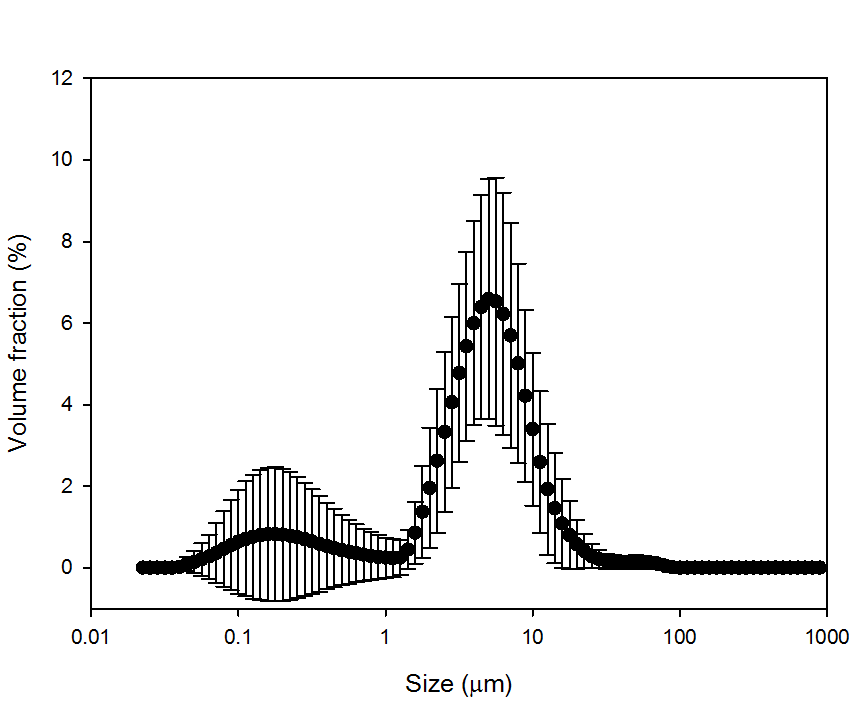

**Supplementary Figure S14.** Volume distribution of PS-*b*-PEO(2000) polymersomes. Molecular weight of PS fragment: 2800-4100. Results as mean ± SD (n = 5).

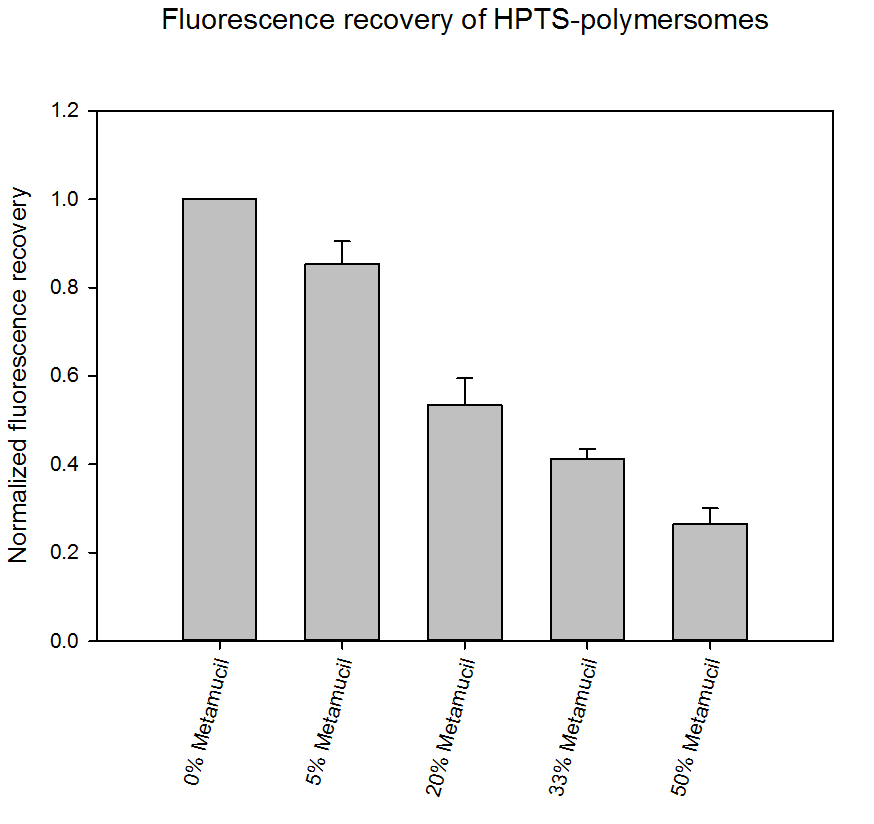

**Supplementary Figure S15.** Normalized fluorescence recovery after incubation of HPTS-containing PS-*b*-PEO polymersomes without transmembrane pH gradient in Metamucil^©^-based hydrogels at pH 6.8. Fluorescence emission intensity normalized to the one at 0% Metamucil^®^ in relation to Metamucil^®^ concentration (0-50%, m/m). HPTS concentration: 30 µM. Inner and outer phase: isotonic phosphate buffer 50 mM at pH 6.8. All results as means ± SD (n = 3).

**Supplementary Table S1.** Diameter of investigated microparticles, liposomes, and polymersomes by laser diffraction (see section 4.3.3).

|  | **D10 (µm)** | **D50 (µm)** | **D100 (µm)** |
| --- | --- | --- | --- |
| Zeolite | 1.5 | 8.4 | 16.6 |
| PMAA | 141 | 235 | 372 |
| Activated charcoal | 0.50 | 0.86 | 1.97 |
| Mesoporous silica | 4.6 | 11.4 | 49.7 |
| DPPC liposomes | 3.1 | 7.3 | 20.4 |
| DSPC liposomes | 3.4 | 7.3 | 16.7 |
| PBD-*b*-PEO polymersomes | 2.2 | 3.9 | 6.9 |
| PS-*b*-PEO polymersomes | 2.1 | 4.1 | 9.3 |

**Supplementary Table S2.** Accuracy and precision of ammonia standard curves of HPTS-loaded PS-*b*-PEO polymersomes with volume fraction of ammonia standard of 17 or 43% (n = 3-5). Values calculated from data in Figure 5A and Supplementary Figure S6.

|  | **Concentration range of ammonia standards** | | | | | |
| --- | --- | --- | --- | --- | --- | --- |
|  | **12.5–200 µM^1^** | | **50–800 µM^1^** | | **12.5–200 µM^2^** | |
| **Ammonia conc. (µM)** | **Accuracy (%)** | **Precision (%)** | **Accuracy (%)** | **Precision (%)** | **Accuracy (%)** | **Precision (%)** |
| 12.5 | 45.4 | 42.6 |  |  | 76.2 | 20.8 |
| 25 | 95.2 | 7.9 |  |  | 95.6 | 11.7 |
| 50 | 107.7 | 3.7 | 69.5 | 24.8 | 102.6 | 0.9 |
| 100 | 109.1 | 2.7 | 88.3 | 16.4 | 105.5 | 7.0 |
| 200 | 97.5 | 0.8 | 107.3 | 4.0 | 98.6 | 1.6 |
| 400 |  |  | 104.7 | 2.7 |  |  |
| 600 |  |  | 101.1 | 2.1 |  |  |
| 800 |  |  | 96.9 | 0.5 |  |  |

^1^Ammonia standard volume fraction: 17%, ^2^ammonia standard volume fraction: 43%

**Supplementary Table S3.** Comparison of the performance parameters of the polymersome assay with reported parameters for other ammonia assays.

| **Assay** | **Coefficient of determination R^2^** | **Lower LOQ (µM)** | **Upper limit (µM)** |
| --- | --- | --- | --- |
| Polymersome assay | 0.9982^a^  0.9932^b^ | 27^a^  39^c^ | 200^a^  800^b^ |
| GLDH-based enzymatic ammonia assay (Randox) | 0.9981^d^ | 123 (manual mode)^d^, 7 (machine-based)[1] | 1180^e^ |
| PocketChem BA | 0.9936^f^ | 52^f^ | 286^e^ |
| Polyaniline nanoparticles[2] | 0.9868 | 36 | 200 |
| Bicompartimental well with ammonia-selective membrane[3] | 0.9757 | 25 | 500 |

^a^12.5–200 µM (43% (v/v); n = 3, Supplementary Figure S6); ^b^12.5–800 µM (17% (v/v); n = 5, Figure 5A); ^c^12.5–200 µM (17% (v/v); n = 5, Figure 5A); ^d^calculated based on Supplementary Figure S7; ^e^according to manufacturer’s instructions; ^f^calculated based on Supplementary Figure S8

**Supplementary Table S4.** Relevant physicochemical properties of the investigated molecules (information retrieved from PubChem, National Center for Biotechnology Information).

| **Molecule** | **Molecular weight (g/mol)** | **pK_a_ of corresponding protonated nitrogen** | **logP** | **Zwitterion at physiological pH** |
| --- | --- | --- | --- | --- |
| Ammonia | 17 | 9.25 | -0.7 |  |
| Dimethylamine | 45 | 10.73 | -0.2 |  |
| Trimethylamine | 59 | 9.80 | 0.3 |  |
| Alanine | 89 | 2.34 (carboxylic acid), 9.60 (amine) | -3.0 | x |
| Glycine | 75 | 2.34 (carboxylic acid), 9.60 (amine) | -3.2 | x |
| Dopamine | 153 | 8.81 (phenol), 10.90 (amine), 13.68 (phenol) | -1.0 |  |
| Propranolol | 259 | 9.42 (amine) | 3.0 |  |
| Levofloxacin | 361 | 6.24 (carboxylic acid), 8.74 (piperazinyl) | -0.4 | x |

**Supplementary Table S5.** Biochemical laboratory parameters of BDL rats of Figure 3D.

|  | **Normal range** | **g^β^** | **h^β^** | **i^β^** | **j^β^** | **k^β^** |
| --- | --- | --- | --- | --- | --- | --- |
| ALT (U/L) | 10-39 | 44 | 57 | 83 | 64 | 84 |
| AST (U/L) | 13-39 | 221 | 307 | 400 | 532 | 619 |
| Albumin (g/L) | 36-45 | 24 | 22 | 20 | 19 | 18 |
| Total bilirubin (µM) | 7-23 | 159 | 197 | 150 | 119 | 137 |
| AP (U/L) | 36-110 | 539 | 634 | 282 | 388 | 587 |
| GGT (U/L) | 9-47 | 27 | 26 | 69 | 44 | 48 |

**Supplementary Table S6.** Properties of PS-*b*-PEO diblock co-polymers purchased from Advanced Polymer Materials.

|  | **Molecular weight, determined by NMR (PS - PEO)^1^** | **Number-average molecular weight (M_n_) in g/mol^1^** | **Dispersity (Đ)^1^** |
| --- | --- | --- | --- |
| PS(1000)-*b*-PEO(2000) | 1040 - 2000 | 3000 | 1.08 |
| PS(1500)-*b*-PEO(2000) | 1560 - 2000 | 3500 | 1.09 |
| PS(2000)-*b*-PEO(2000) | 1970 - 2000 | 4000 | 1.07 |
| PS(2500)-*b*-PEO(2000) | 2600 - 2000  2770 - 2000 | 4600  4700 | 1.09  1.09 |
| PS(3000)-*b*-PEO(2000) | 3150 - 2000 | 5000 | 1.19 |
| PS(3500)-*b*-PEO(2000) | 3570 - 2000 | 6500 | 1.13 |
| PS(4000)-*b*-PEO(2000) | 3900 - 2000 | 6000 | 1.15 |
| PS(5000)-*b*-PEO(2000) | 5150 - 2060 | 7200 | 1.07 |
| PS(6000)-*b*-PEO(2000) | 6000 - 2180 | 8200 | 1.09 |

^1^According to manufacturer

**Supplementary Table S7.** Properties of synthesized PS-*b*-PEO diblock co-polymers.

| **Polymer batch ID** | **Molecular weight, determined by NMR (PS - PEO)^1^** | **Number-average molecular weight (M_n_)^1^** | **Dispersity (Đ)** |
| --- | --- | --- | --- |
| F2 | 5500 - 2000 | 7400 | 1.45 |
| F3 | 3900 - 2000 | 6900 | 1.45 |
| F4 | 4300 - 2000 | 7200 | 1.53 |
| FF1 | 4500 - 2000 | 7500 | 1.45 |
| FF2 | 4500 - 2000 | 7200 | 1.55 |
| N1 | 2800 - 2000 | 7200 | 1.48 |
| N2 | 3800 - 2000 | 6500 | 1.43 |
| N3 | 4300 - 2000 | 6300 | 1.43 |
| N5 | 3300 - 2000 | 5900 | 1.38 |
| N6 | 4100 - 2000 | 7200 | 1.44 |
| A16 | 3300 - 2000 | 6300 | 1.40 |

**References**

[1]	I. Seiden-Long, K. Schnabl, W. Skoropadyk, N. Lennon, A. McKeage, Evaluation of a third party enzymatic ammonia method for use on the Roche Cobas 6000 (c501) automated platform, Clin. Biochem. 47 (2014) 1116–1120. doi:10.1016/J.CLINBIOCHEM.2014.04.022.

[2]	N.T. Brannelly, A.J. Killard, An electrochemical sensor device for measuring blood ammonia at the point of care, Talanta. 167 (2017) 296–301. doi:10.1016/J.TALANTA.2017.02.025.

[3]	O.B. Ayyub, A.M. Behrens, B.T. Heligman, M.E. Natoli, J.J. Ayoub, G. Cunningham, et al., Simple and inexpensive quantification of ammonia in whole blood, Mol. Genet. Metab. 115 (2015) 95–100. doi:10.1016/J.YMGME.2015.04.004.
